# Target binding triggers hierarchical phosphorylation of human Argonaute-2 to promote target release

**DOI:** 10.1101/2022.01.06.475261

**Authors:** Brianna Bibel, Elad Elkayam, Steve Silletti, Elizabeth A Komives, Leemor Joshua-Tor

## Abstract

Argonaute (Ago) proteins play a central role in post-transcriptional gene regulation through RNA interference (RNAi). Agos bind small RNAs (sRNAs) including small interfering RNAs (siRNAs) and microRNAs (miRNAs) to form the functional core of the RNA Induced Silencing Complex (RISC). The sRNA is used as a guide to target mRNAs containing either partially or fully complementary sequences, ultimately leading to down regulation of the corresponding proteins. It was previously shown that the kinase CK1α phosphorylates a cluster of residues in the eukaryotic insertion (EI) of Ago, leading to the alleviation of miRNA-mediated repression through an undetermined mechanism. We show that binding of miRNA-loaded human Ago2 to target RNA with complementarity to the seed and 3’ supplemental regions of the miRNA primes the EI for hierarchical phosphorylation by CK1α. The added negative charges electrostatically promote target release, freeing Ago to seek out additional targets once it is dephosphorylated. The high conservation of potential phosphosites in the EI suggests that such a regulatory strategy may be a shared mechanism for regulating miRNA-mediated repression.

## Introduction

Most metazoan genes are regulated post-transcriptionally by RNA interference (RNAi)^1^. This is a conserved process whereby an Argonaute (Ago) protein binds to a small RNA, ∼22 nucleotides (nt) in length, called a microRNA (miRNA), and uses it as a guide to seek out and bind target mRNAs containing regions of partial complementarity, typically in the target’s 3’UTR of genes^1,2^. Binding of the core RNA Induced Silencing Complex (RISC), consisting of Ago and an miRNA guide^3^, to an mRNA can lead to recruitment of cofactors and decreased translation of protein from that mRNA through a number of mechanisms including: preventing translation initiation and elongation; promoting mRNA decay; triggering premature termination and co-translational protein degradation; and sequestering targets from the translational machinery^4,5^. Ago has a high affinity for miRNAs with a very slow off rate^6,7^. Consequently, most Ago/guide complexes are long lived, with average half-lives on the order of days^8^ and some complexes stable for at least 3 weeks^9^. Thus, once loaded with a guide, Ago can repress countless matching mRNA sites, as long as it is able to efficiently release these targets once silencing is achieved. This turnover is crucial because potential target sites are present in large excess compared to the corresponding miRNAs^10,11^.

miRNA-binding sites are typically located within the target mRNA’s 3’ UTR and are primarily determined by perfect complementarity in a 6-8 nt “seed sequence” corresponding to nucleotides 2-8 of the guide miRNA (g2-8)^12^, augmented by additional pairing beyond this region. In the core RISC, the subseed region of the miRNA (g2-5) is preorganized for base-pairing^13–15^, which allows Ago to rapidly sample potential target sites^15^. Binding to a fully-complementary seed induces conformational changes in Ago that expose more of the miRNA and widen Ago’s RNA-binding channel, allowing for further pairing^16^. Target sites vary in their amount of complementarity to a guide outside of the seed sequence, and additional pairing can influence a site’s repression efficacy^17^. This pairing often occurs in the more solvent-exposed 3’ supplemental region (corresponding to g12-17) through a second nucleation event^2,18^. 3’ supplemental pairing, most effectively with 3-4 contiguous base pairs centered around g13-16^17^, can strengthen repression by increasing guide/target affinity through a decrease in *K*_*off*_^19–21^. It also allows miRNAs of the same family (miRNAs that share the same seed sequence) to preferentially repress different sets of targets^22,23^.

Ago proteins are part of a larger Argonaute superfamily whose members show high structural conservation throughout evolution and have 4 main structural domains: N, PAZ, MID, and PIWI^24^. The MID and PIWI domains form an almost continuous lobe which, together with the N domain, forms a crescent-shaped base above which the PAZ domain is able to move more freely^24,25^. The 5’ end of the small RNA is bound by a pocket created by the MID and PIWI domains^26^ and the 3’ end is held by the PAZ domain^27,28^, with the length of the small RNA cradled in a groove traversing the crescent^13,14,29^. The PAZ domain is the most flexible and, sometimes in concert with the N domain, undergoes rigid body movements along linker regions (L1 & L2) in order to accommodate RNA^16,18,25,30^. Humans have four Ago orthologs (hAgo1-4) in addition to four Piwi-clade Agos, with hAgo2 being the most highly expressed in most tissues^31,32^. Although particular functions have been attributed to specific hAgo proteins, hAgo1-4 are largely considered functionally redundant^33^. The lone exception is hAgo2; knock-out of hAgo2 is lethal in early mouse development^34,35^, whereas knock-out of the other hAgos, and even knock-out of all three, is not^36^. This essentiality is attributed largely to the fact that, under most circumstances, only hAgo2 has “slicer” activity – the ability to cleave fully-complementary targets between nucleotides across from g10 and g11^24,34,37^. Although slicing is not required for miRNA-mediated regulation, it is necessary for the maturation of rare but important Dicer-independent miRNAs including miR-451^38^ and miR-486^39^, which play essential roles in erythrocyte development.

Although the core architecture is conserved among all Argonaute proteins, they are subject to multiple types of post-translational modification, including ubiquitinylation, PARylation, and phosphorylation. For example, S837 undergoes stress-induced p38/MAPK-mediated S837 phosphorylation, which, among other effects, is thought to contribute to hAgo2 localization in processing bodies (P-bodies)^40^. There is a span of residues on the surface of the PIWI domain in eukaryotic miRNA-specialized Ago proteins which is absent in prokaryotic Ago proteins as well as siRNA- or piRNA-specialized Argonaute superfamily proteins (e.g *Drosophila melanogaster* Ago2 and Piwi proteins, respectively). This region, corresponding to residues ∼820-837 in hAgo2 numbering, is termed the Eukaryotic Insertion (EI), and it contains 4-5 highly conserved potential phosphorylation sites; four serine residues (in hAgo2 numbering, S824, S828, S831, and S834) as well as a threonine (T830) which is only conserved in Ago2 homologs. This cluster was recently found to be heterogeneously phosphorylated in multiple Ago homologs in a variety of cell types and organisms^41,42^ and its phosphorylation and dephosphorylation were attributed to the kinase CK1α and the phosphatase PP6/ANKRD52, respectively^41^.

This cycle of phosphorylation of Ago’s EI was implicated in regulatory control of Ago itself^41,42^. Golden et al. found that *decreased* EI phosphorylation led to an increase in target RNA association with the Ago-guide complex in cell culture, as measured by qt-RT-PCR of immunoprecipitated hAgo2, whereas *increased* EI phosphorylation or its mimicking had the opposite effect^41^. Additionally, disruption of the hAgo EI phosphorylation cycle led to dysregulation of mRNA levels, as determined by RNA-seq. In the absence of EI phosphorylation, hAgo2 associated with a more diverse pool of mRNA targets, but with decreased coverage of individual targets as determined by CHIP-Seq^41^. Furthermore, reporter assays showed that increased phosphorylation or its mimicking led to reduced repression of reporter targets in cell culture^38^ and in *C. elegans*^39^. These silencing defects could be rescued by tethering Ago to the target, indicating that phosphorylation impaired a step upstream of repression, preventing productive Ago-guide/target interactions^41,42^. Together with their findings that miRNA production and guide association appeared unaffected, these results point to a role of EI phosphorylation in mediating Ago-guide/target interactions, but key questions remained about the molecular events associated with this process.

Using a combination of biochemical and biophysical assays, we determined that binding of miRNA-bound hAgo2 to targets containing 3’ supplemental pairing triggers hierarchical phosphorylation of the EI by CK1α. The added negative charge electrostatically repels the negatively-charged target RNA, leading to target release. Such regulation could help explain how cells maintain tight and consistent miRNA-mediated regulation despite there being a large excess of miRNA target sites compared to Ago and miRNA molecules^10,11^.

## Results

### Target binding promotes CK1α-mediated EI phosphorylation in vitro

Previous work showed that hAgo2 EI phosphorylation was enhanced by the addition of a miRNA accompanied by a miRNA sponge containing multiple binding sites specific to that miRNA^41^. This suggested that, within cells, EI phosphorylation is stimulated by target binding. This stimulatory effect could be due to features inherent to the ternary Ago-miRNA/target complex itself, or caused by binding of external mediators, e.g. repression-mediating proteins that are recruited after target binding such as GW-182, PAN2-PAN3, and CCR4-NOT ^4^. To determine if target-binding alone could stimulate phosphorylation in the absence of cofactors, we tested CK1α-mediated phosphorylation of recombinant, dephosphorylated, hAgo2 in its apo state (RNA-free), when bound to a guide miRNA (miR-200b), and when bound to guide and a partially-complementary target *in vitro*. For this target, we used a 30-nt RNA we will refer to as ZT1, which corresponds to a natural, highly-conserved, miR-200 binding site in the 3’ UTR of the transcription factor ZEB1^43^. We did not detect any phosphorylation of RNA-free hAgo2 above background and detected very low levels of phosphorylation with guide-only-bound hAgo2, 0.14 ± 0.07 mol phosphorylation per mol Ago2 (Figure 1). However, when we added target RNA, we observed substantial phosphorylation, 2.6 ± 0.2 mol phosphorylation per mol hAgo2 (Figure 1). No phosphorylation was observed when all 5 phosphorylatable EI residues were mutated to alanine hAgo2(5XA) (Figure 1), confirming that the observed phosphorylation of the wild-type (wt) hAgo2 occurs and is dependent upon EI residues. To test whether the other human Ago orthologs behave in a similar fashion, we tested hAgo1 and hAgo3 using a this assay. We found that these Agos are also subject to target-binding-triggered CK1α-mediated phosphorylation *in vitro* (Figure 1 – figure supplement 1). This is consistent with previous findings that hAgo1 and hAgo3 are phosphorylated on multiple residues in the EI *in vivo*^42^.

**Figure 1:**
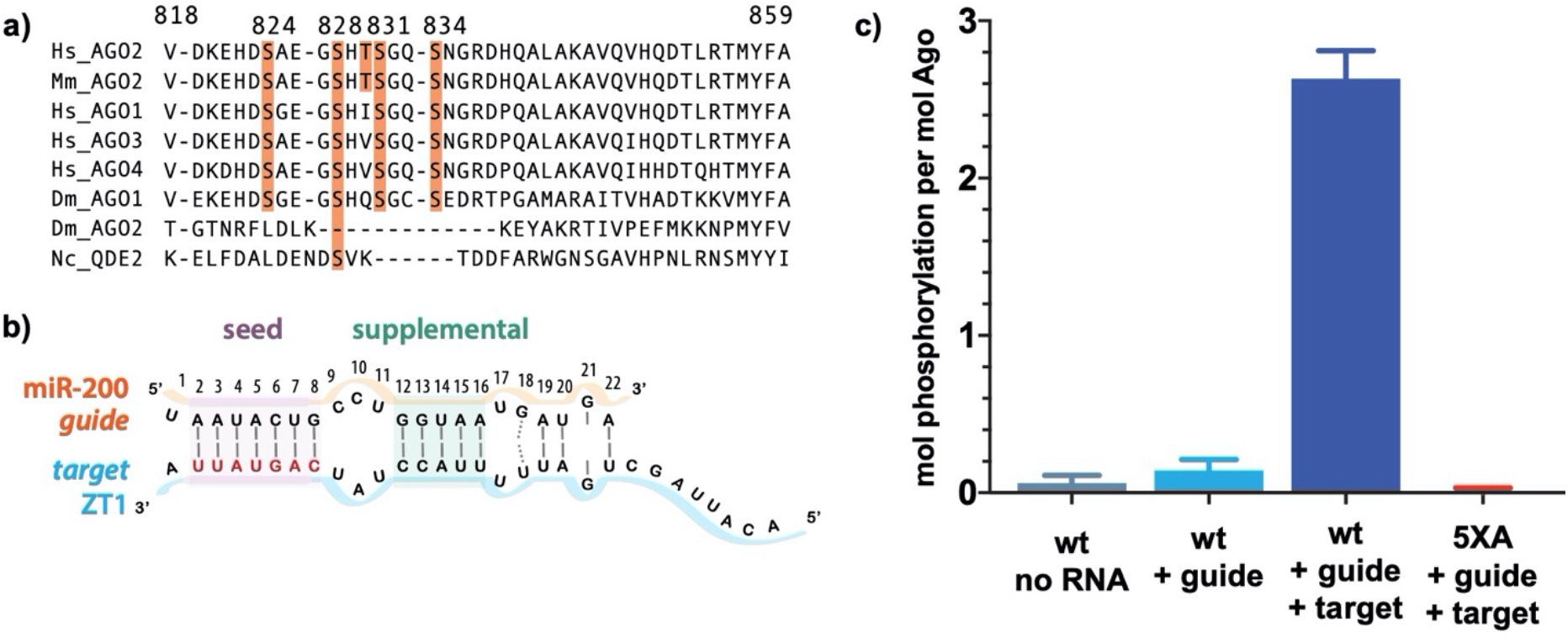
Target binding triggers phosphorylation of the hAgo2 Eukaryotic Insertion (EI). a) Conservation of the EI in eukaryotic miRNA-handling Argonaute proteins. Phosphorylatable residues are highlighted in orange. The 4 serines are highly conserved in higher eukaryotes, whereas the threonine is specific to hAgo2. Residue numbering is based on hAgo2. Hs, *Homo sapiens*; Ce, *Caenorhabditis elegans*; Dm, *Drosophila melanogaster*; Mm, *mus musculus*. b) Guide/target pair corresponding to mirR-200 and a 30-mer target RNA (ZT1) corresponding to a natural and validated miR-200 binding site located in the 3’UTR of ZEBT1. Nucleotides of the target complementary to the miR-200 seed sequence are labeled in red and the seed and 3’ supplementary pairing regions are indicated. c) Target binding, but not guide binding, dramatically increases CK1a-mediated phosphorylation of hAgo2, as measured by in vitro phosphorylation assays using the guide/target pair shown in b. For this and all other in vitro phosphorylation assay figures, unless otherwise noted, reactions were carried out in triplicate and measured using liquid scintillation counting. Quantification of phosphorylation was done based on spotted ATP and plotted ± standard error (SE) as described in Materials and Methods.

### Phosphorylation is promoted by pairing in the supplemental region

Because of the observed effects of target binding, we decided to further investigate what features of the target RNA affect EI phosphorylation. In order to determine the minimum target length sufficient to trigger it, we performed *in vitro* phosphorylation assays with a series of ZT1 target RNAs, starting with an 8-nt target complementary to the seed region and extending the 5’ end (corresponding to the 3’ end of the guide) up to a target length of 30-nt. No significant phosphorylation was detectable up to a target length of 13-nt, however, phosphophorylation increased dramatically once the target length reached 14-nt, with further increases in phosphorylation observed at longer lengths (Figure 2a). These differences could not be explained by differences in binding affinity (Figure 2 – figure supplement 1).

**Figure 2:**
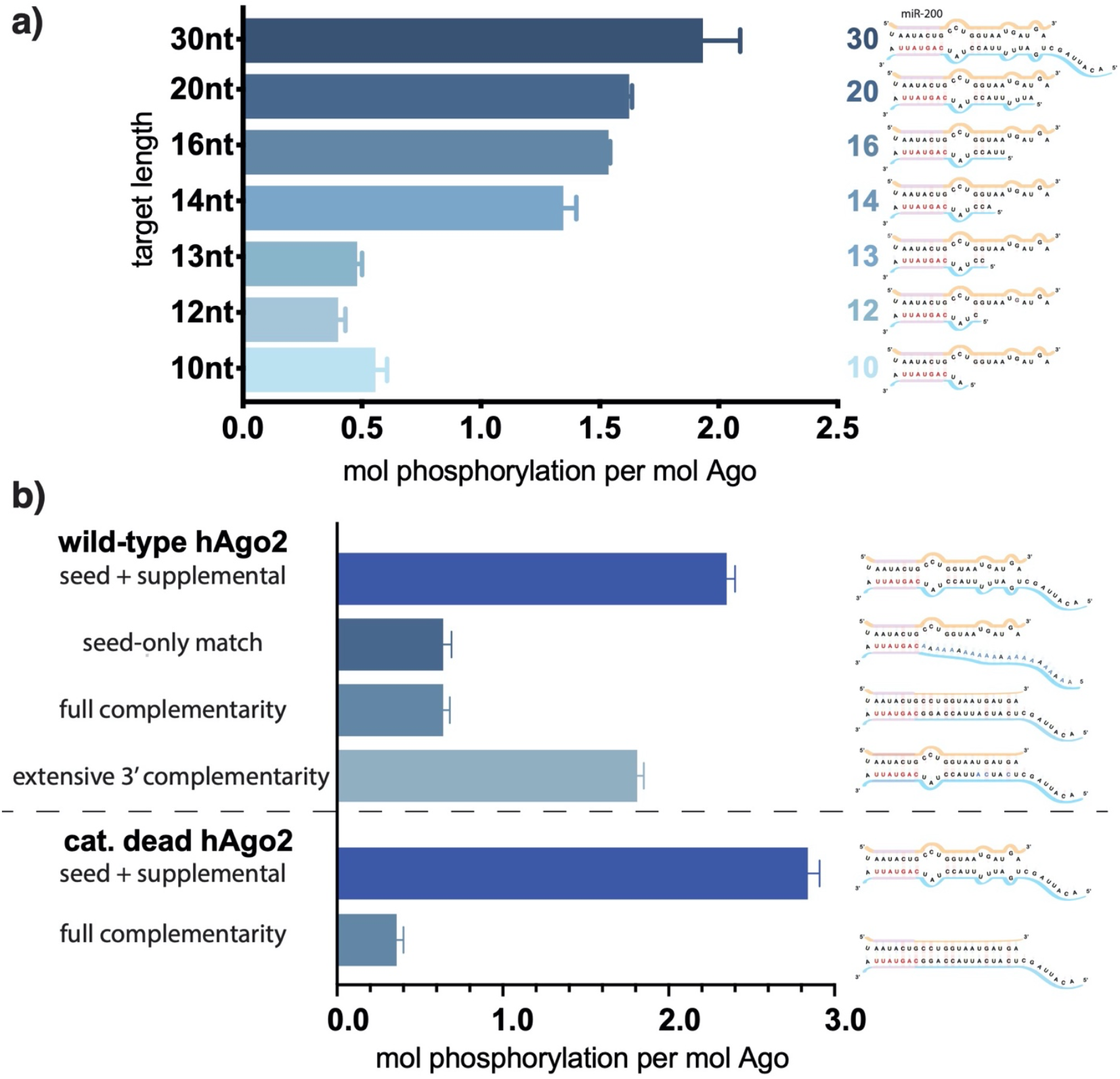
Target length and complementarity requirements. a) In vitro phosphorylation assays with ZT1-based targets of varying lengths show that a minimum of 14 nucleotides is needed to trigger robust phosphorylation. b) Assays with ZT1-based targets with different levels of complementarity to the guide (miR-200) show that the seed + 3’ supplemental configuration is best able to induce phosphorylation. A catalytically-inactivated (cat-dead) hAgo2, hAgo2(669N) is included to control for possible slicing of the fully-complementary target. Quantification of phosphorylation was done based on spotted ATP and plotted ± SE as described in Materials and Methods. n=3

It is likely that 14 nt represents the shortest length of target capable of establishing stable pairing in the supplemental region. Therefore, to further investigate the importance of supplemental pairing, we tested phosphorylation of hAgo2-guide-traget ternary complexes with varying amounts of complementarity to the guide miRNA. Length-matched (30 nt) target with complementarity to the seed region alone was incapable of stimulating robust hAgo2 phosphorylation (Figure 2b). However, targets with full complementarity also failed to trigger phosphorylation in either wild-type or catalytically-inactivated hAgo2 (Ago2(D669N))^34^. The latter was included to control for potential confounding effects caused by slicing, which occurs with hAgo2 and fully-complementary guide/target pairs. This result is consistent with our finding that EI phosphomimetics do not significantly affect slicing (Figure 2 – figure supplement 2). Furthermore, targets with extended 3’ complementarity, resembling target-directed miRNA decay (TDMD) targets, which are known to lead to an altered conformation of hAgo2^44^, triggered significantly less phosphorylation than the original target (Figure 2b). Therefore, length and supplemental region complementarity are not sufficient to maximally trigger phosphorylation. Instead, a more particular target site configuration of seed+supplemental pairing (i.e., without central or extensive 3’ pairing) serves as a trigger.

These findings may be explained by the unique conformations Ago adopts to accommodate varying amounts of base pairing. Binding to short targets is known to lead to a shift in hAgo2’s PAZ domain^16^, which could make the EI more accessible for CK1α. There are no structures of hAgo2 bound to a full-length “canonical” miRNA target containing seed and supplemental pairing. However, the crystal structure of a catalytically-inactivated hAgo2 bound to a target RNA with complementary to nucleotides 2-16 of the guide (g2-g16) was recently determined^18^. In this structure, likely due to Ago’s inability to cleave the target, the RNA is not base paired in the central region (g9-11) and instead pairs only in the seed and supplemental regions, a conformation which the authors posit could represent one that is adopted by Ago upon binding traditional miRNAs^18^. In this structure, the 3’ end remains held by the PAZ domain while the N & PAZ domains move away from the MID & PIWI domains, which could make the EI more accessible to CK1α.

### Triggering of phosphorylation upon target binding is not due to major conformational changes in the EI

The EI itself is unresolved in all Ago structures determined to date. In order to obtain structural information regarding conformational changes that the EI might be undergoing during these events, we turned to hydrogen-deuterium exchange mass spectrometry (HDX-MS), which measures conformational dynamics and solvent accessibility by analyzing the exchange of hydrogens for deuterium over time. The measured deuterium uptake corresponds primarily to exchange from backbone amides, and thus regions with stable secondary structure have high levels of protection (low deuterium uptake) whereas more flexible and/or exposed regions are less protected and thus have higher deuterium uptake^45,46^. Samples of hAgo2 in RNA-free (RF), guide-bound (G-bound), and guide+target-bound (GT-bound) states were analyzed to investigate whether guide- and/or target-binding influence the structure of the EI or the surrounding residues. Since no reliable peptides covering the EI could be measured for wild-type hAgo2, we used the 5XA mutant, which has a similar target-binding affinity compared to dephosphorylated wild-type (Figure 3 – figure supplement 1), yet did have reliable coverage of the EI. The 5XA mutant showed minimal differences in exchange compared to the dephosphorylated wild-type RNA-free hAgo2 over the rest of the protein (Figure 3 – figure supplement 2). Very minor, yet statistically-significant (p<0.01), differences were only seen at one of the four time points (2 min) and were restricted to 10 (out of 255) peptides, none of which were in the region surrounding the EI. Thus, 5XA is a good proxy for the dephosphorylated wild-type protein. Near complete (99.4%) sequence coverage was achieved, with a total of 262 peptides and an average redundancy of 5.29 (Figure 3 – figure supplement 3).

Seven peptides covering the EI were detected in all datasets, the shortest being 811-842 and 812-842. These peptides were consistent with each other, as would be expected given the almost complete overlap between the two. Importantly, they showed no statistically-significant differences between the G-bound and the GT-bound at any timepoint (p<0.01) (Figure 3). The G-bound showed slight protection compared to the RF, however taking overlapping peptides into account to increase resolution, this increase can be attributed to higher uptake in the region downstream of 834, outside of the EI itself (Figure 3a). The EI showed a moderately high level of exchange, suggesting little defined secondary structure in any state, and no detectable differences in secondary structure content between the states. If maximal exchange was already realized at the earliest timepoint (30s), we would not be able to detect an increase in flexibility nor solvent accessibility, only a decrease. Since we do not see any such decrease, we can conclude that neither guide-nor target-binding offer protection of phosphorylatable residues. However, we cannot rule out conformational changes occurring that do not involve strong changes in secondary structure content.

**Figure 3:**
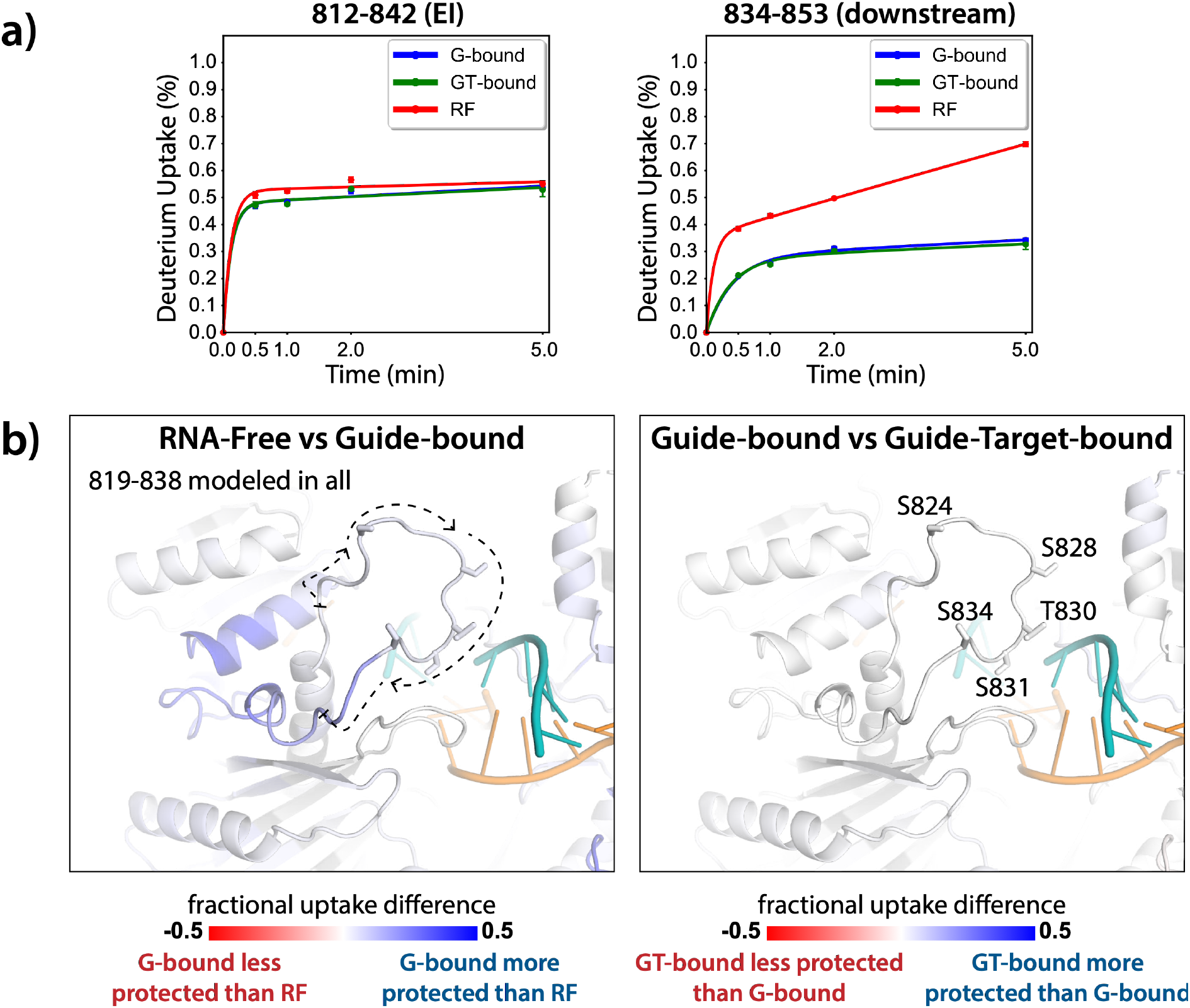
HDX-MS of the hAgo2(5XA) EI. a) Uptake plots for the shortest peptide covering with the EI shows no significant difference between the guide-bound (G) and guide+target-bound (GT) states. The increased uptake in the RNA-free (RF) state can be accounted for by increased uptake in region downstream of the phosphorylation sites. b) Fractional uptake differences between the RF and G-bound (left) or the G-bound and GT-bound hAgo2(5XA) (right). Residues 819-838 were modeled into the structure of hAgo2 bound to a target with seed and supplementary pairing (6n4o) in Chimera using MODELLER’s loop refine tool. Filtered data for the 30 s time point are mapped onto the model structure and phosphorylatable residues are displayed in stick representation and labeled.

The lack of differences between the RF and G-bound states within the EI is in stark contrast to the large differences observed outside of the EI. The most substantial protection upon guide-binding occurred in regions known to interact directly with the miRNA, such as the 5’ binding pocket (Figure 4 – figure supplement 1). However, the RF also showed higher relative uptake over most of the protein, suggesting greater overall flexibility prior to guide-binding. This is consistent with previous results showing guide-binding protects from limited proteolysis^13^ and with the observation that RNA-free hAgo2 is unstable in cells and rapidly degraded^47^, whereas guide-bound hAgo2 complexes are incredibly stable^9^. Differences between the G-bound and GT-bound states were more subtle. Significant differences were localized mainly to regions expected based on crystal structures, including the L1 linker and helix 7 of the L2 linker region (Figure 4), which undergo a rigid body movement upon target binding, widening the N-PAZ channel to accommodate pairing beyond the seed sequence^16^.

**Figure 4:**
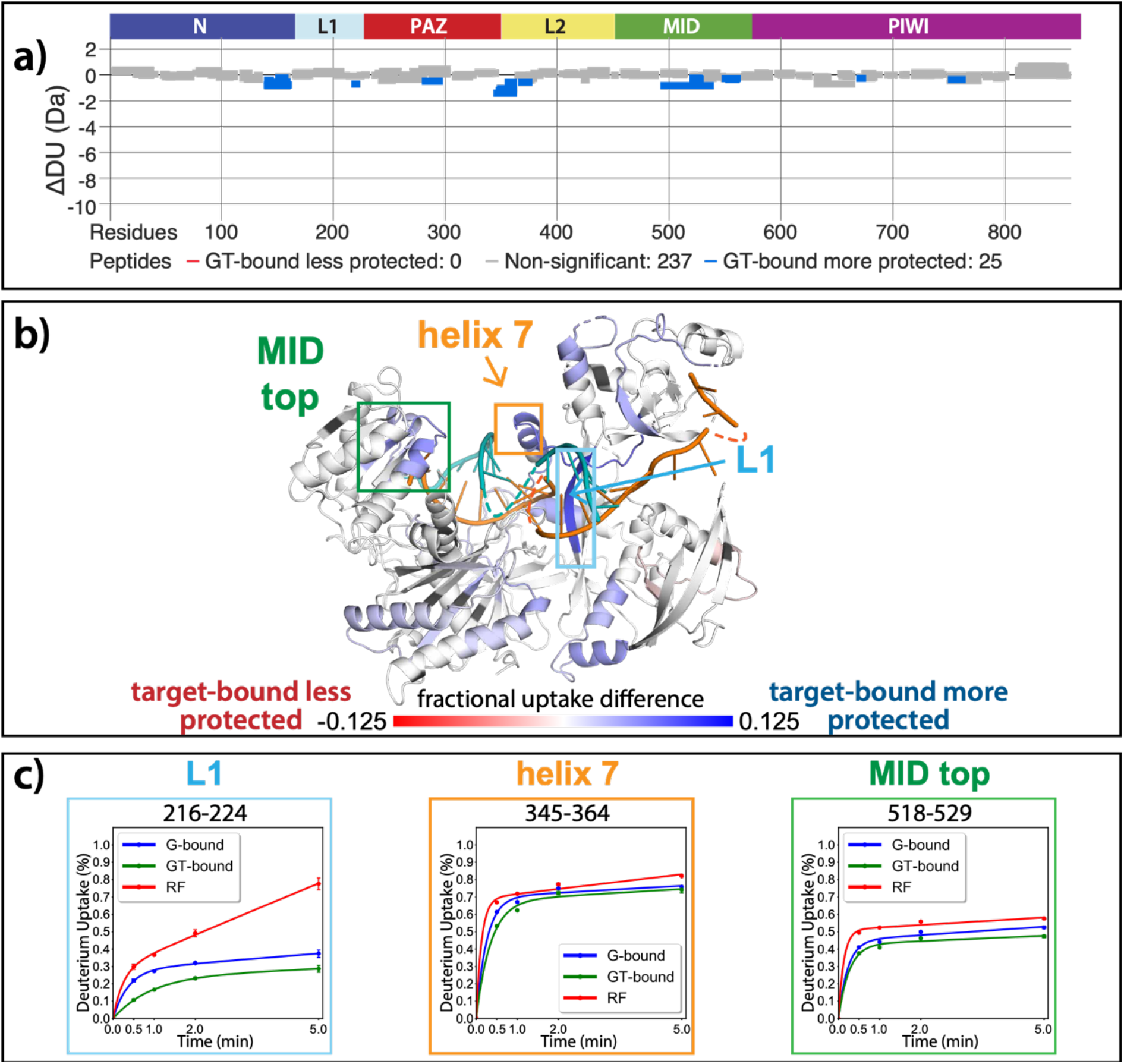
Effects of target-binding on deuterium exchange are structurally concentrated and small in magnitude. a) Wood’s Differential plot comparing the absolute deuterium uptake of guide-bound 5XA hAgo2 (G-bound) and guide+target-bound 5XA hAgo2 (GT-bound) at the 30 second time point. Statistically-different (p <0.1) peptides are highlighted in blue indicating that the GT-bound is somewhat more protected than the G-bound. Positions of hAgo2 domains are displayed above. b) Fractional uptake difference comparing G-bound and GT-bound 5XA hAgo2 at the 30s time point. Statistically-different peptides were used to generate a heat map displayed on the structure of hAgo2 bound to a guide and target with seed and supplementary pairing (PDB code 6N4O, Sheu-Gruttadauria et al. (2019)), scaled to the data range (blue where target is 12.5% more protected to 12.5% where target is 12.5% less protected). c) Deuterium uptake plots for peptides representative of three main areas showing protection offered upon target binding.

### Phosphorylation occurs hierarchically from S828

We next sought to determine where in the EI phosphorylation was occurring. CK1α has a canonical sequence preference of pS/pT-x-x-S/T* or (E,D)_n_-x-x-S/T*-x where “p” indicates a phosphorylated site, * indicates the site to be phosphorylated, and “x” indicates any amino acid^48^. Thus, prior phosphorylation (by CK1α or another kinase) can prime for subsequent phosphorylation, making CK1α capable of hierarchical phosphorylation. This has been proposed to occur at the EI^41^, however heterogeneity has hindered past attempts to characterize EI phosphorylation status at the amino-acid level. In addition to differences in phosphorylation status among Ago molecules, identical charged states (e.g. 1P, 2P, 3P, or 4P) can be formed with different combinations of phosphorylated residues. Furthermore, due to the close clustering of the sites, as well as the tendency of phosphorylations to be lost during processing and mass spectrometry, proteomic studies have been unable to identify every phosphorylation combination^41,42^.

In order to determine which EI residues get phosphorylated *in vitro*, we addressed this ambiguity by performing *in vitro* phosphorylation assays on hAgo2 constructs in which we prevented phosphorylation at specific sites by mutating each potentially phosphorylatable serine or threonine to alanine, individually or in various combinations. Since we had previously determined that target-binding is required for phosphorylation (Figure 1), we carried out these assays in the presence of guide (miR-200) and target (ZT1). We started by looking at which residues could get phosphorylated by CK1α *in vitro* without priming phosphorylation by way of creating a series of five “single o” hAgo2 mutations in which only a single residue is “open” for phosphorylation, with the others changed to alanine. For example, to check for S824 phosphorylation, we designed an Ago2(S828A/T830A/S831A/S834A) mutant, abbreviated to “824o”. We observed phosphorylation of the 828o mutant, but not of any of the other four (Figure 5a). Thus, only S828 can get phosphorylated by CK1α *in vitro* in the absence of prior priming phosphorylation. An upstream acidic cluster in the EI was proposed to provide a non-primed canonical recognition sequence^41^; however, we found that phosphorylation was not impacted when we mutated those acidic residues to neutral ones (i.e. E821Q/D823N/E826Q) (Figure 5 – figure supplement 1).

**Figure 5:**
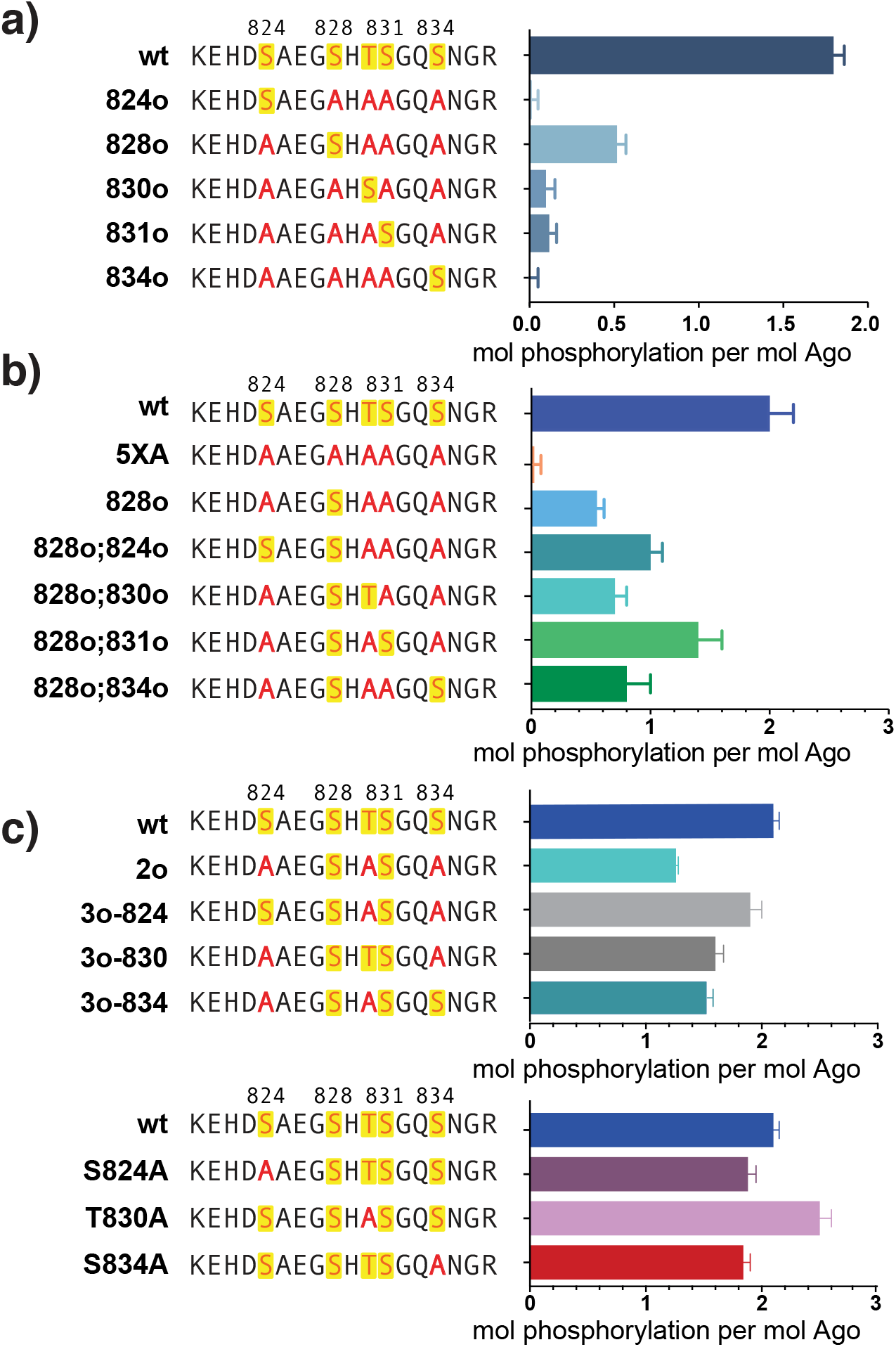
Phosphorylation of the hAgo2 EI occurs hierarchically. a) CK1α-mediated phosphorylation of miR-200/ZT1-bound wt, 5XA, and “single open” mutants in which only a single potential phosphosite is open for phosphorylation by CK1α, with the rest mutated to alanine, shows that S828 is the only site capable of CK1α-mediated phosphorylation without previous priming phosphorylation. b) Determining priming from pS828. “Double open” mutants in which only two sites are open for phosphorylation, S828 and one other site, show that S828 is capable of priming for S831 phosphorylation. c) Determining priming from pS831. “Triple open” mutants in which only three sites are open for phosphorylation, S828, S831, and S834, show that S831 is capable of priming for further phosphorylation, but that the third site phosphorylation is more heterogeneous (top), as supported by similar effects of single alanine substitutions (bottom). Quantification of phosphorylation was done based on spotted ATP and plotted ± standard error (SE) as described in Materials and Methods.

To examine the possibility that CK1α-mediated phosphorylation of S828 may serve a priming role for additional EI phosphorylations, we first tried a S828 phosphomimetic (S828E). This failed to prime for further phosphorylation (Figure 5 – figure supplement 2), however glutamate is different in size and charge to phosphoserine and thus may not sufficiently recapitulate the effects of true phosphorylation^49^. Therefore, we engineered “double o” hAgo2 EI constructs in which 828 and only one other residue are open for phosphorylation, while the other 3 sites are mutated to alanine. In these constructs, S828 becomes phosphorylated and can potentially serve to prime further phosphorylation of the other available site, resulting in increased phosphorylation compared to the singly phosphorylatable 828o mutant. The “double o” mutant in which S828 and S831 were open (i.e. Ago2(T830A, S831A, S834A)) showed significantly increased phosphorylation, whereas the mutants with 824, 830, or 834 open showed only minor increases (Figure 5b). This suggested that pS828 is capable of priming for phosphorylation of S831.

To determine whether pS831 is subsequently capable of priming for further phosphorylation, we created “triple o” mutants: 824o;828o;831o [Ago2(T830A;S834A)]; 828o;831o;834o [Ago2(S824A;T830A)]; and 828o;830o;831o [Ago2(S824A;S834A)]. We saw an increase in phosphorylation in all three cases, suggesting that, after the second phosphorylation at S831, the EI phosphorylation pattern can follow different routes and is more heterogenous than the initial S828 followed by S831 phosphorylation pattern (Figure 5c). Individual mutation of S824 or S834 to alanine slightly reduced phosphorylation, and to similar extent, wheras mutation of T830 to alanine actually increased phosphorylation compared to wt. The increase in phosphorylation seen in the T830A mutation may indicate that this mutation makes the other sites better substrates, potentially by decreasing steric hindrance.

### Phosphorylation promotes target release

Next we wanted to study the functional consequences of this phosphorylation on the interaction of minimal RISC (hAgo2+guide) with target RNA. Because, as shown above, phosphorylation can only be achieved *after* target binding, we conducted target release assays in which we phosphorylated hAgo2 bound to the guide RNA and radiolabeled target RNA (ZT1). We then used a timecourse filter-binding assay to measure release after dilution into a 250-fold excess of unlabeled target. CK1α-mediated phosphorylation of wt hAgo2 significantly increased *k*_*off*_ by ∼7.5-fold, and thus decreased the half-life of the RISC/target complex (Figure 6a). These effects are specific to EI phosphorylation, as CK1α-mediated treatment of hAgo2(5XA) had no effect. These effects are additive – the more sites available for phosphorylation, the larger the observed effect (828o < 828o;831o < 828o;831o;834o) (Figure 6b). The “triple o” mutant 824o;828o;831o showed similar effects as 828o;831o;834o (Figure 6 – figure supplement 1) suggesting that the overall charge, rather than the specific phosphorylated sites is most important for target release.

**Figure 6:**
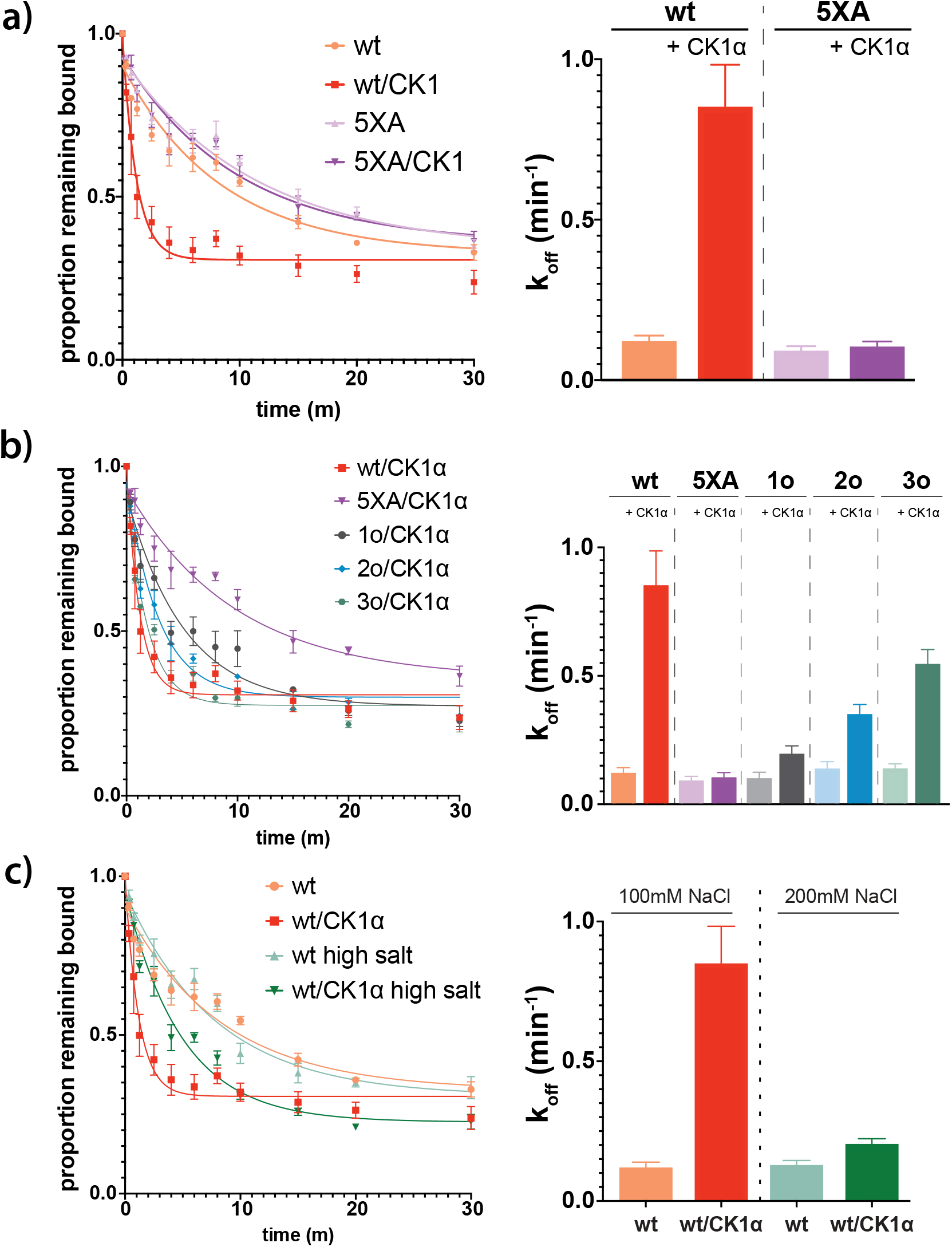
Phosphorylation promotes target release. a) Filter-binding based target release assay tracking the release of labeled target from wild-type (wt) hAgo2 phosphorylated while bound to unlabeled target. b) Target release assays performed as in a with 1 open, 2 open, and 3 open (824o;828o;831o) hAgo2 mutants show that increasing phosphorylation decreases half life. c) The effects of phosphorylation on target release are minimal under high salt conditions (200 mM NaCl vs 100 mM NaCl), showing that the effect on target release is largely electrostatic. a-c) Proportion of target remaining bound over time since addition of an unlabeled target chase is plotted with 95%CI (left) and half lives calculated using a One-Phase Exponential Decay equation in GraphPad Prism are shown ± SE (right). n=4 for 3o-834/CK1a, n=3 for others.

It was proposed that the negative charge of the added phosphates could clash with the negative charge of the target RNA backbone, leading to loosening of hAgo2 binding to the target by charge repulsion^42^. To test this, we repeated the release assay with high salt concentrations, where the high ionic strength should provide electrostatic shielding, thus minimizing charge-based effects. In these conditions, differences between the off rate of CK1α-treated and untreated wt hAgo2 were much smaller, only ∼1.5-fold higher for phosphorylated hAgo2, supporting the hypothesis that the effects of phosphorylation on promoting target release are largely electrostatically mediated (Figure 6c).

## Discussion

miRNA-mediated gene repression is a widespread mode of regulation in metazoans. Most human genes contain at least one evolutionarily-conserved miRNA target site, and many genes are under regulation by multiple families of miRNAs. Each miRNA family, in turn, typically has hundreds of targets; the 90 most broadly conserved miRNA families have over 400 conserved targets each^1^. At the same time, the number of miRNA target sites is much greater than the number of miRNA-Ago complexes^10^ and the half-lives of mRNAs are, on average, significantly shorter than those of RISC complexes^7^. Therefore, it appears that each core RISC targets multiple mRNAs throughout its lifetime. On the other hand, repression efficacy is largely related to binding stability, with stronger binding correlating with enhanced repression^50^. Thus, ensuring robust mRNA repression at the global level is a balancing act between enabling strong binding at individual sites and binding to a large number of sites. 3’-supplemental pairing increases binding affinity and thus can lead to enhanced repression at cognate sites^51^, but high affinity may come with the risk of effectively sequestering RISC. CK1α-mediated EI phosphorylation provides a mechanism to counterbalance this added affinity, freeing RISC to repress additional targets. Indeed, Golden et. al found that hAgo2 which could not be phosphorylated in the EI associated with a more diverse pool of mRNA targets, but with decreased coverage of individual targets, accompanied by global dysregulation of miRNA-mediated repression^41^.

This may help explain discrepancies between the *in vitro* and *in vivo* effects of 3’ supplemental pairing. *In vitro*, 3’-supplemental pairing can significantly enhance the stability of Ago/target interactions, with 3-4 contiguous canonical base pairs in the supplemental region increasing affinity over 20-fold for some miRNA/target pairs^18,19,21^. This effect is more pronounced for miRNAs with weak, low GC-content, seeds. However, the contributions of supplemental pairing *in vivo* and in cell culture have been reported to be less substantial^17,20,21^. Some of this discrepancy has been attributed to differences in cell types and measurement conditions, as well as the large effects of a small number of miRNA/targets being diluted out in large datasets^19^. However, as we have shown in this study, supplemental pairing triggers phosphorylation of the EI would offset the added stability. Therefore, *in vivo*, the effects of supplemental pairing on target repression would be expected to be less pronounced.

CK1α phosphorylation likely occurs in a distributive manner^52^. Because phosphorylation of multiple residues would require CK1α to actively interact with Ago multiple times, supplemental pairing might be expected to increase phosphorylation by way of added affinity. However, we found that target retention time is not the primary driver of CK1-α mediated EI phosphorylation, as demonstrated by the minimal levels of phosphorylation observed upon binding to the fully-complementary or TDMD-like target. Our findings instead suggest that the conformation of the RISC complex is of primary importance with regards to triggering phosphorylation. Compared to 3’ supplemental targets, fully-complementary targets require different conformational changes due to base pairing of the central region^18^, and these changes might make Ago’s EI a poorer substrate for CK1α.

Recently, a study linked a variety of germline single amino acid mutations in hAgo2 to RNAi deficits and neurological symptoms^53^. These mutations clustered in helix 7, L1, and a loop in the PIWI domain, and, when expressed in cell culture, some of the mutated Agos had decreased EI phosphorylation and a slight increase in quantity of select target mRNAs. This, combined with molecular dynamics simulations, led the authors to propose that reduced phosphorylation prevents efficient target release. Based on our findings that target-binding promotes phosphorylation, prolonged target binding in the usual conformation would be expected to increase rather than decrease phosphorylation. Therefore, it appears possible that the identified mutations may cause hAgo2 to adopt a suboptimal target-bound conformation which disfavors, or at least fails to promote, phosphorylation.

There are multiple examples in the literature where substrate tertiary structure has been shown to be critical for promoting CK1-mediated phosphorylation. For example, T-antigen is phosphorylated by CK1 at non-canonical sites at its N-terminus in a manner that is dependent on proper folding and an intact C-terminus^54^. Similarly, conformational changes induced by target binding may cause Ago to adopt a conformation which is structurally preferable for CK1α. However, this preference cannot soley be attributed to the changes making the EI more accessible. Despite the high degree of flexibility and solvent accessibility of the EI and the surrounding area in the RNA-free hAgo2, as seen from our HDX-MS results (Figure 3 and Figure 4 – figure supplement 1) and through limited proteolysis^13^, we were unable to phosphorylate it *in-vitro* (Figure 1). This suggests there must be an additional factor required. Since the EI shows no significant changes in deuterium exchange neither upon guide-nor upon target-binding, the additional information needed to enable phosphorylation stems from outside of the EI. Such information could come from a different part of the hAgo2 protein that is now in a more suitable conformation to interact with the kinase. Alternatively and/or additionally, factors outside of hAgo2 may assume this role, potentially the target RNA itself.

As we show in this study, target-binding triggers the CK1α-mediated phosphorylation of S828 (Figure 1). However, this residue is a noncanonical target for CK1α, deviating from its preferred consensus sequence, as determined largely through peptide-based studies. Numerous examples have now come to light in which CK1α phosphorylates non-canonical sites, and additional consensus sequences have been identified^55^. Nevertheless, CK1 kinases are thought to generally prefer primed substrates. Binding of the phosphorylated, priming residue, or a cluster of acidic residues to a basic patch on the kinase, named Anion Binding Site 1, is thought to stabilize the kinase in an active conformation and position the downstream phosphorylation site in the active site^52,56,57^. However, phosphorylation was not impacted when we neutralized the acidic residues previously proposed to serve as a priming cluster^41^ (Figure 5 – figure supplement 1). We would like to suggest that it is the target RNA’s backbone, being negatively-charged, that may be able to bind to this site and prime for S828 phosphorylation. The need for seed+supplementary pairing as opposed to other complementarity combinations in order to trigger robust phosphorylation supports a role for conformational change. These changes might position the RNA to interact with the kinase properly, rather than making the EI more accessible.

Because miRNA-mediated repression does not involve slicing of the targets, additional cofactors are required to carry out silencing of miRNA targets. After miRNA-bound Ago binds a target mRNA, Ago binds a scaffolding protein of the GW182 family (TNRC6A/B/C in humans) which in turn recruits deadenylation complexes (PAN2-PAN3 and/or CCR4-NOT) and decapping complexes to promote target degradation^4^. Previous studies have found that EI phosphorylation does not impact hAgo2/GW182 interaction^41,42^. It is possible that one of the recruited proteins recruits CK1α. However, since we are able to observe Ago phosphorylation in the absence of additional proteins, they are not required for CKα-mediated phosphorylation. Importantly, because we did not observe phosphorylation until after target binding, and even then, can only detect initial phosphorylation of a single site, the *in vitro* system appears to recapitulate a regulated system having high specificity.

Because Ago must be dephosphorylated by PP6 before it can optimally repress additional targets, phosphorylation also has the potential to serve as a global regulatory measure. Knocking out PP6 in human cell lines led to a global decrease in RISC binding to mRNA^41^. Under normal conditions, however, CK1α-mediated EI phosphorylation is rapidly countered by dephosphorylation^41^. Neither 5XA nor 5XE hAgo2 could rescue miRNA-mediated repression in Ago^-/-^cells, indicating that the phosphorylation cycling process itself is important, not merely the presence or absence of phosphorylation.

Our findings that *only* S828 can be phosphorylated in our *in vitro* system without priming phosphorylation, and that this phosphorylation only occurs after binding to targets meeting certain criteria reveal novel structural and sequence determinants of highly specific CK1α-mediated phosphorylation. We therefore propose that target binding serves as a means for regulating CK1α-mediated phosphorylation of Ago, which, in turn, regulates RISC turnover and thus miRNA-mediated mRNA repression. The presence of multiple phosphorylation sites with decreasing target affinity may merely reflect the biophysical requirements for establishing sufficient negative charge to repel the target RNA. However, it might also provide a way for a cell to titrate the Ago/target affinity and/or to serve as a “checkpoint” in order to ensure that targets aren’t spuriously released.

This mechanism appears specialized for repression mediated by miRNA, as opposed to siRNA, as binding to fully-complementary sites only triggers minimal phosphorylation (Figure 2b) and phosphomimicking does not impair slicing (Figure 2 – figure supplement 2). This is consistent with the evolutionary conservation of the phosphosites among miRNA-specialized Ago proteins (e.g. *Drosophila melanogaster* Ago1 (dmAgo1)) but not among siRNA-specialized Agos (e.g. dmAgo2), and our work provides a potential mechanistic explanation. With siRNA, slicing of the fully-complementary target promotes its release by destabilizing the guide/target duplex^21^, but partially-complementary miRNAs with high affinity, such as those with 3’ supplemental pairing, could get stuck on a target and phosphorylation could serve as a mechanism to promote their release in the absence of slicing. Though we showed that hAgo1 and hAgo3 are also subject to target-binding-triggered CK1α-mediated phosphorylation *in vitro* similar to hAgo2, we did not investigate the functional impact of this phosphorylation; however, based on the high homology between the human Ago’s and their apparent redundancy in most instances^33^, we would expect the effects of hAgo1 and hAgo3 phosphorylation to be similar to those we observed with hAgo2. Therefore, this regulatory mechanism is likely shared by these miRNA-handling Ago orthologs, suggesting a potentially broad impact on post-transcriptional regulation.

In conclusion, our findings show that Ago EI phosphorylation is a post-target-binding regulatory mechanism that promotes target release, freeing guide-bound Ago, namely RISC, from the mRNA target and dissuading it from binding more targets until it gets dephosphorylated by PPP6C/ANKRD52^41^. This dephosphorylation was previously shown to be rapid^41^, therefore the released Ago-guide complex is likely to quickly be able to seek out and repress additional targets. As long as the cycle of phosphorylation/dephosphorylation is not disrupted, this would allow cells to strike a balance between the increased efficacy and specificity provided by stronger binding sites and the quest for efficiency, allowing a single miRISC to repress multiple mRNAs (Figure 7).

**Figure 7:**
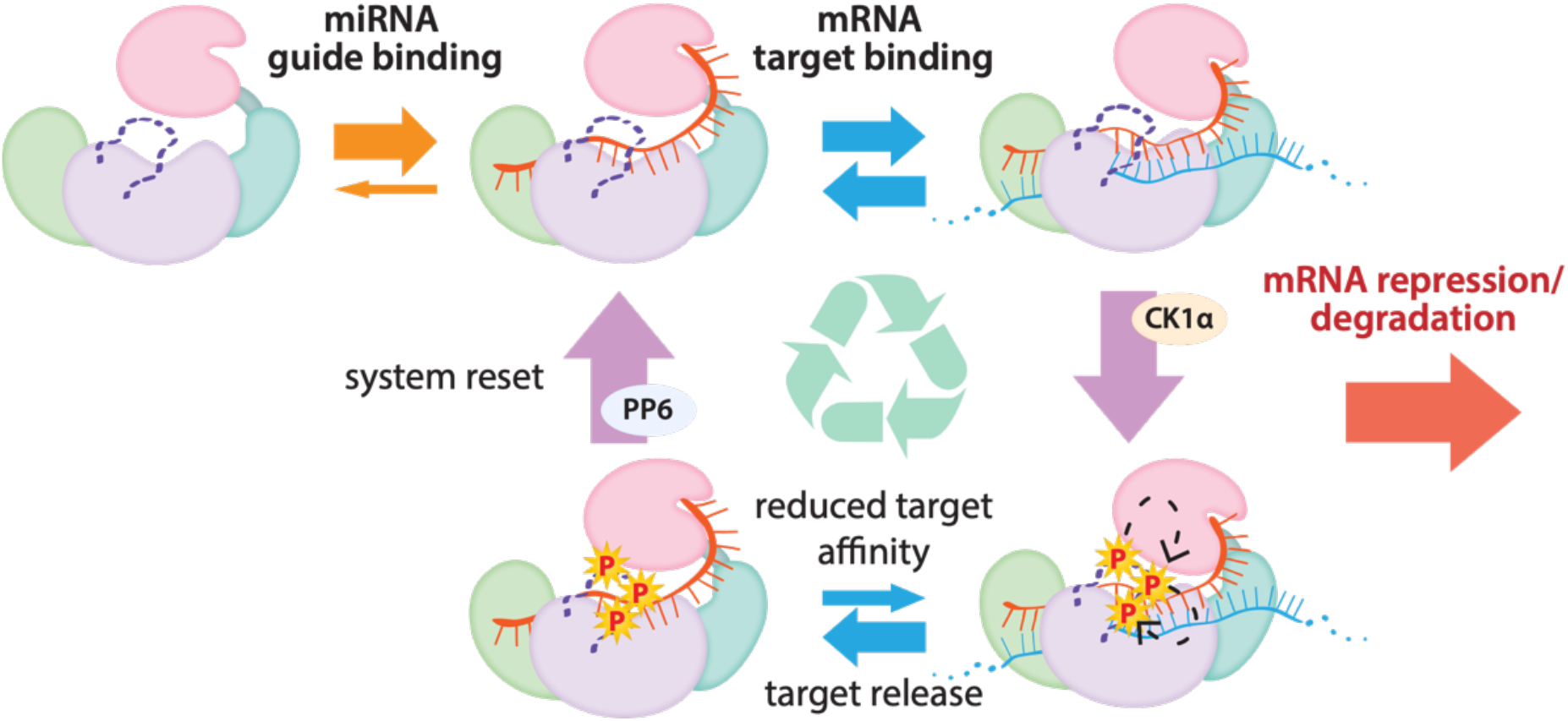
The cycle of phosphorylation/dephosphorylation plays a regulatory role in silencing. In order to repress a target mRNA, Ago binds to a small RNA guide to form the core RISC complex, which subsequently binds to target mRNAs containing sequence complementarity to the guide to effect repression. The core RISC complex (Ago/guide) is incredibly stable but would have to release target mRNA prior to repression of additional targets. We propose that this release is facilitated by CK1α-mediated phosphorylation of the EI. We showed that phosphorylation is triggered by binding to targets that are not fully complementary, but that contain complementarity to the seed and 3’ supplementary pairing regions of the guide. Binding to such targets results in the complex adopting a productive conformation for EI phosphorylation. We suggest this may be due to orienting of the RNA backbone phosphates to serve as noncanonical priming phosphates. EI phosphorylation occurs hierarchically starting at S828 and the additive negative charges of the phosphates electrostatically repel the target RNA, leading to target release. The phosphorylated RISC has reduced affinity for further target binding. However, the system is quickly reset by dephosphorylation. This allows the RISC complex to bind to and repress additional sites, helping explain how RISC can function efficiently and effectively in the context of excess target sites.

This mechanism of promoting miRNA target release appears specialized for miRNA-mediated repression and the broad conservation of the phosphorylation sites suggests that its use is widespread among miRNA-handling Ago proteins.

## Acknowledgments

We thank all members of the Joshua-Tor lab for helpful suggestions and comments. We also thank Nicholas Tonks (Cold Spring Harbor Laboratory) for valuable discussions and use of equipment. Additional thanks to Sheena D’Arcy and Naifu Zhang (University of Texas at Dallas) for helpful discussions. We are grateful to Julian Langer and Jonathan Zoller (Max Plank Institute of Biophysics) for sharing an R script for statistically-filtering peptides and demonstrating how to use it. We thank the CSHL Mass Spectrometry Shared Resource, which is supported by Cancer Center Support Grant 5P30CA045508. BB was supported by the NSF Graduate Research Fellowship Program and CSHL School of Biological Sciences. LJ is an investigator of the Howard Hughes Medical Institute. The HDX-MS was performed on the Synapt G2Si HDX mass spectrometer obtained by shared instrumentation grant S10 OD016234.

## Materials and Methods

### Protein expression and purification

Recombinant hAgo2 was prepared as previously described,^13^ with the addition of a dephosphorylation step. Briefly, hAgo2 was expressed in Sf9 insect cells with an N-terminal StrepII_SUMO fusion tag using the MultiBac system^58^. After Strep-Tactin affinity chromatography (Strep-Tactin Superflow High Capacity, IBA) (50 mM Tris pH 8, 100 mM KCl, 5 mM DTT) the tag was removed using TEV protease and RNA-free Ago was separated from Ago loaded with endogenous (insect) RNAs by cation exchange chromatography with a stepwise KCl gradient elution (MonoS 10/100, Cytiva). RNA-free Ago was treated overnight at 4°C with λ protein phosphatase (λPP) at a 1:2.5 molar ratio, with MnCl_2_ added to a final concentration of 1 mM. The protein was then further purified via size exclusion chromatography (Superdex 200 10/300 Increase GL, Cytiva)(10 mM Tris pH 8, 200 mM KCl, 5 mM DTT). Aliquots were frozen at -80°C, 2 mg/mL, 10% glycerol. hAgo1 and hAgo3 were prepared in a similar manner.

Human CK1α (isoform 1) was expressed in Sf9 insect cells with an N-terminal StrepII_SUMO fusion tag using the MultiBac system. Following Strep-Tactin affinity chromatography (50 mM Tris pH 8, 200 mM NaCl, 5 mM DTT), the tag was removed using TEV protease and CK1α was further purified by cation exchange chromatography (MonoS, linear NaCl gradient elution) and size exclusion chromatography (Superdex75 Increase 10/300 GL, Cytiva)(10 mM Tris pH 8, 200 mM NaCl, 5 mM DTT). Aliquots were frozen at -80°C, 0.5 mg/mL, 50% glycerol.

λPP was cloned into a pET28 vector with an N-terminal His tag and expressed in BL21(DE3) *E. coli*. It was induced with 1 mM IPTG and expressed overnight at 18°C in TB media. It was then purified by Ni-NTA affinity chromatography (Qiagen)(50mM Tris pH 8, 300mM NaCl, 25 mM imidazole, 0.5mM TCEP, 0.1 mM MnCl_2_; eluted with 250 mM imidazole) followed by size exclusion chromatography (Superdex75)(50 mM Tris pH 8, 150 mM NaCl, 0.5 mM TCEP, 0.1 mM MnCl_2_). Activity was verified using a colorimetric pNPP assay and aliquots were frozen at -80°C in 50% glycerol, 2.7 mg/mL.

### In vitro phosphorylation assays

In vitro phosphorylation was carried out with hAgo2:guide:target at 1 μM and CK1α at 20 nM. Prior to adding CK1, Ago was incubated 1:1 with guide RNA for 30 minutes, followed by incubation with target RNA (1:1:1 molar ratio) for 30 minutes, both steps were carried at room temperature. The complex was then mixed with kinase reaction buffer (final concentrations of 200μM total ATP, 2 nM [γ-^32^P]-ATP (Perkin-Elmer), 25 mM Tris pH 7.4, 10 mM MgCl_2_, 2.5 mM DTT, 0.5 mM Na_3_VO_4_). CK1α was then added and reactions were incubated at 37°C for 90 min at which time they were quenched with a stop buffer containing 2 5mM EDTA and 25 mM unlabeled ATP. Aliquots were spotted on phosphocellulose paper, washed three times (5 min each) in 75 mM phosphoric acid, one time in acetone, and then air dried and measured using liquid scintillation counting. Moles of phosphate per aliquot were calculated by comparison to spotted (unwashed) reaction buffer. This value was divided by the number of moles of Ago in the aliquot to determine the mole phosphorylation detected per mole Ago.

### Target binding

Equilibrium target binding assays were performed using a double-membrane filter-binding setup with a slot blot vacuum filtration device (BioRad SF) containing an upper nitrocellulose membrane (to capture protein and protein/RNA complexes) and a lower nylon membrane (to capture free RNA) as previously described. Briefly, hAgo2 and guide were mixed at a 1:1 molar ratio and incubated at room temperature for 30 minutes, then serially diluted in 10 mM Tris pH 8, 100 mM NaCl, 5 mM DTT and added to 5’ ^32^P-labeled target RNA (50-100 pM). After 1h incubation to come to equilibrium, protein (including RNA-bound protein) and free RNA were separated by slot blot. Results were obtained through phosphorimaging (Typhoon FLA 7000, Cytiva) and quantified using GeneTools (SynGene). Kinetic parameters were fit using GraphPad Prism (Version 9), one site specific binding.

For competition target binding assyas, hAgo2 was mixed 1:1 with miR-200 and allowed to bind for 30 min at RT. This RISC complex was then added to a serial dilution mixture of radiolabeled ZT1 (30-nt), which was held constant at 50 pm, and unlabeled competitor target which was 1:3 serially diluted, starting at 750 nM for the seed-only complementary target (ZT1.som) and 500 nM for all others. The mixture was allowed to equilibrate for 1 hour at RT before separating protein from free RNA using a filter-binding assay as above. Data was quantified in GeneTools (SynGene) and Ki was fit using GraphPad Prism.

### Guide binding

hAgo2 and guide were mixed at a 1:1 molar ratio and incubated at room temperature for 60 minutes, then serially diluted in 10mM Tris pH 8, 100 mM NaCl, 5mM DTT and added to 5’ ^32^P-labeled miR-200 (25 pM). After 1h incubation to reach equilibrium, protein and protein-RNA complex were separated from free RNA by slot blot and quantified as above.

### Target release

hAgo2 was mixed 1:1 with miR-200 guide RNA and incubated for 30 min at RT. A mix of unlabeled and ^32^P-labeled ZT1 target RNA (30 nt) was added and incubated for a further 30 min. *In vitro* phosphorylation was then carried out with 500 nM hAgo2/guide, 200 nM labeled ZT1, 300 nM unlabeled ZT1 and 20 nM CK1α for 60 min at 37°C followed by 30 min at 10°C. An aliquot was then diluted with a high concentration of unlabeled target to a final concentration of 5 nM hAgo2, 2 nM labeled ZT1, 500 nM unlabeled ZT1, in either standard buffer (10 mM Tris pH 8.0, 100 mM NaCl, 5 mM DTT) or high salt buffer (10 mM Tris pH 8.0, 200 mM NaCl, 5 mM DTT) at 10°C. An initial sample was taken immediately after mixing (at 15 s) and set as the “0” timepoint. Aliquots were taken out to 30 min and proportions bound were compared to the proportion bound in the “0” sample. At each timepoint, free and protein-bound RNA were separated via slot blot, followed by autoradiography and quantification using GeneTools. Data were fit to a One-Phase Exponential Decay equation using GraphPad Prism and plotted ± SE. Reactions carried out in triplicate.

### Slicer assays

Slicer assays were carried out as previously described with slight modifications.^30^ Briefly, reactions were performed in 10 mM Tris-HCl (pH 8.0), 100 mM KCl, 10 mM DTT, 2 mM MgCl2, and 10% glycerol. hAgo2 protein was loaded with at miR-200 at a 1:1 ratio and incubated for 30 min at at RT. 20 μl slicing reactions were initiated by mixing 5’ ^32^P-labeled target RNA (2 nM final concentration) in the reaction buffer with 8 nM loaded hAgo2:miR20a complex. After 60 min, the reaction quenched in formamide loading buffer and incubated at 95°C for 5 min. The radiolabeled slicing products were separated using urea PAGE, visualized by phosphorimaging, and quantified with GeneTools.

### HDX-MS

HDX-MS was carried out by the HDX Core of the University of California, San Diego Biomolecular/Proteomics Mass Spectrometry Facility (UCSD BPMSF) (NIH shared instrumentation grant S10 OD016234). Hydrogen/deuterium exchange mass spectrometry (HDXMS) was performed using a Waters Synapt G2Si equipped with nanoACQUITY UPLC system with H/DX technology and a LEAP autosampler. Individual proteins were purified by size exclusion chromatography in 10 mM Tris pH 8, 200 mM KCl, 5 mM DTT, 10% glycerol immediately prior to analysis. The final concentrations of proteins in each sample were 5 μM. For each deuteration time, 4 μL complex was equilibrated to 25 °C for 5 min and then mixed with 56 μL D_2_O buffer (10 mM Tris pH 8, 200 mM KCl, 5mM DTT, 10% glycerol in D_2_O) for 0, 0.5, 1, 2, or 5 min. The exchange was quenched with an equal volume of quench solution (3 M guanidine hydrochloride, pH 2.66).

The quenched sample (50 μL) was injected into the sample loop, followed by digestion on an in-line pepsin column (immobilized pepsin, Pierce, Inc.) at 15°C. The resulting peptides were captured on a BEH C18 Vanguard pre-column, separated by analytical chromatography (Acquity UPLC BEH C18, 1.7 μM, 1.0 × 50 mm, Waters Corporation) using a 7-85% acetonitrile gradient in 0.1% formic acid over 7.5 min, and electrosprayed into the Waters SYNAPT G2Si quadrupole time-of-flight mass spectrometer. The mass spectrometer was set to collect data in the Mobility, ESI+ mode; mass acquisition range of 200–2,000 (m/z); scan time 0.4 s. Continuous lock mass correction was accomplished with infusion of leuenkephalin (m/z = 556.277) every 30 s (mass accuracy of 1 ppm for calibration standard). For peptide identification, the mass spectrometer was set to collect data in MS^E^, ESI+ mode instead.

The peptides were identified from triplicate MS^E^ analyses of 10 μM solution of protein in (10 mM Tris pH 8, 200 mM KCl, 5 mM DTT, 10% glycerol), and data were analyzed using PLGS 3.0 (Waters Corporation). Peptide masses were identified using a minimum number of 250 ion counts for low energy peptides and 50 ion counts for their fragment ions. The peptides identified in PLGS were then analyzed in DynamX 3.0 (Waters Corporation) and, for displaying uptake, the deuterium uptake was corrected for back-exchange as previously described^59^ using DECA Software (https://github.com/komiveslab/DECA). The relative deuterium uptake for each peptide was calculated by comparing the centroids of the mass envelopes of the deuterated samples vs. the undeuterated controls following previously published methods^61^. The experiments were performed in triplicate, and independent replicates of the triplicate experiment were performed to verify the results (Supplemental table 1).

Wood’s differential plots were created using 2.0^62^, with peptides prior to back-exchange correction. For creating PyMol heat maps showing regions with significantly different deuterium exchange, peptides were filtered for significant differences in dynamic uptake curves using a custom R script, provided by Dr. Julian Langer, which employs a two-stage t test based on Houde et al., 2011^63^ and described in Eisinger et al., 2017^64^. Briefly, in the first stage, the mean deuterium uptake ± SEM for each peptide was subjected to a t test (n=3, p≤0.05, two-sided, unpaired) and peptides that passed for at least three timepoints were taken to the second stage. In this stage, the summed differences for the peptides were subjected to a second t test (n=4, p≤0.01, two-sided, unpaired) to test for overall uptake differences in the dynamic uptake curve. Peptides that passed both filters were used to generate a heat map in DECA^59^ that was mapped onto structural models of hAgo2 in PyMol (Schrödinger, LLC).

### RNAs used (synthesized by Dharmacon)

miR-200 (hsa-miR-200b-3p): 5’-pUAAUACUGCCUGGUAAUGAUGA-3’ validation of miR-200 ZT1 sites: https://www.nature.com/articles/ncb1722

ZT1 (miR-200 binding site corresponds to position 1243-1250 of ZEB1 3’UTR): 5’-ACAUUAGCUGAUUUUUACCUAUCAGUAUUA-3’

ZT1e3c (extended 3’ complementarity): 5’-ACAUUAGCUCAUCAUUACCUAUCAGUAUUA-3’

ZT1s (sliceable (fully complementary)): 5’-ACAUUAGCUCAUCAUUACCAGGCAGUAUUA-3’

ZT1som (seed-only match): 5’-AAAAAAAAAAAAAAAAAAAAAACAGUAUUA-3’

ZT1.20: 5’-AUUUUUACCUAUCAGUAUUA-3’

ZT1.16: 5’-UUACCUAUCAGUAUUA-3’

ZT1.14: 5’-ACCUAUCAGUAUUA-3’

ZT1.13: 5’-CCUAUCAGUAUUA-3’

ZT1.12: 5’-CUAUCAGUAUUA-3’

ZT1.11: 5’-UAUCAGUAUUA-3’

ZT1.10: 5’-AUCAGUAUUA-3’

---

**Figure 1 – figure supplement 1:**
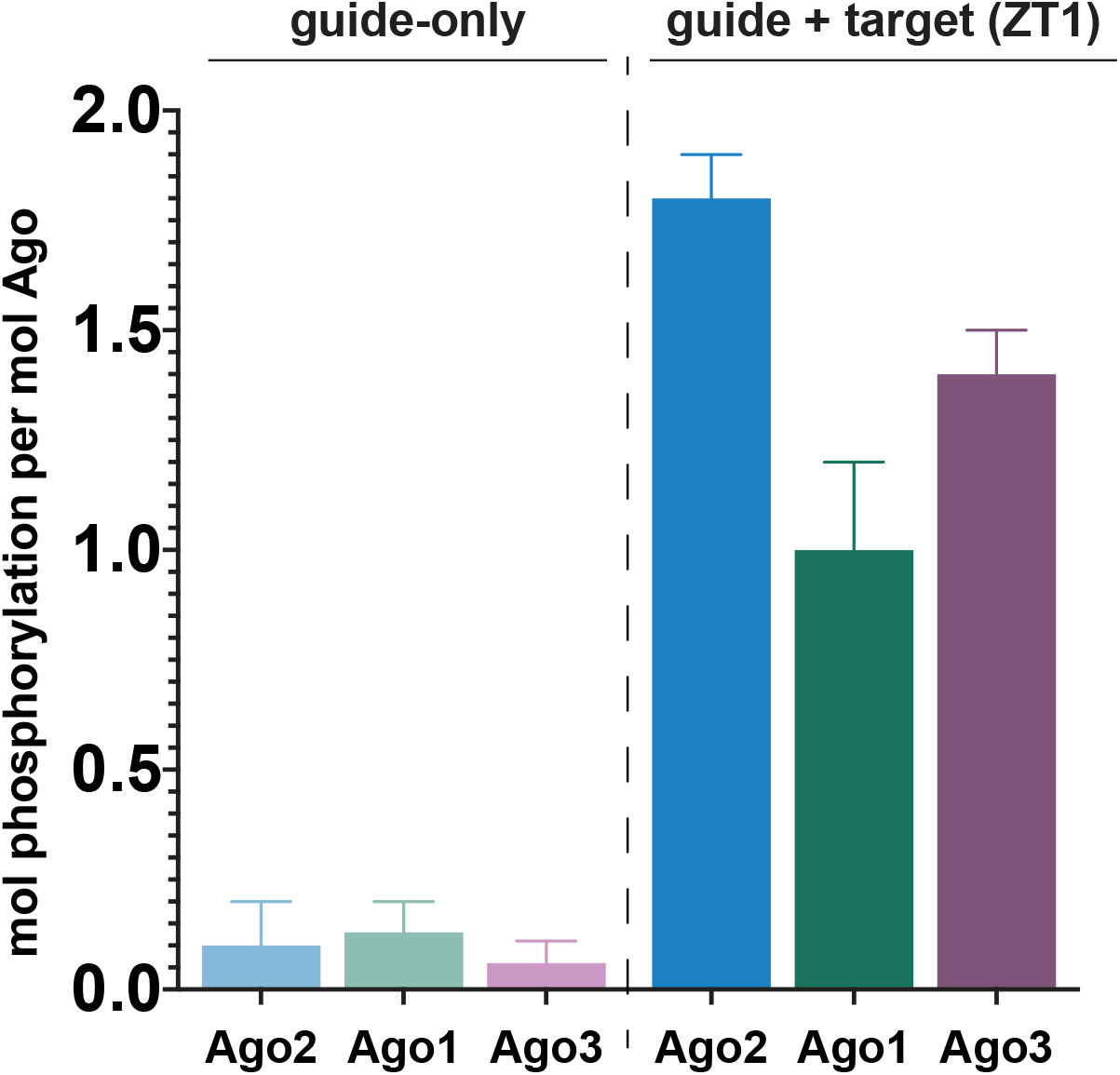
Target binding triggers phosphorylation of the EI in hAgo1 & hAgo3. In vitro phosphorylation assays of human Agos show that hAgo1-3 all undergo target-triggered phosphorylation. hAgo4 was not tested due to difficulty in its expression. Quantification and graphing as in Figure 1.

**Figure 2 – figure supplement 1:**
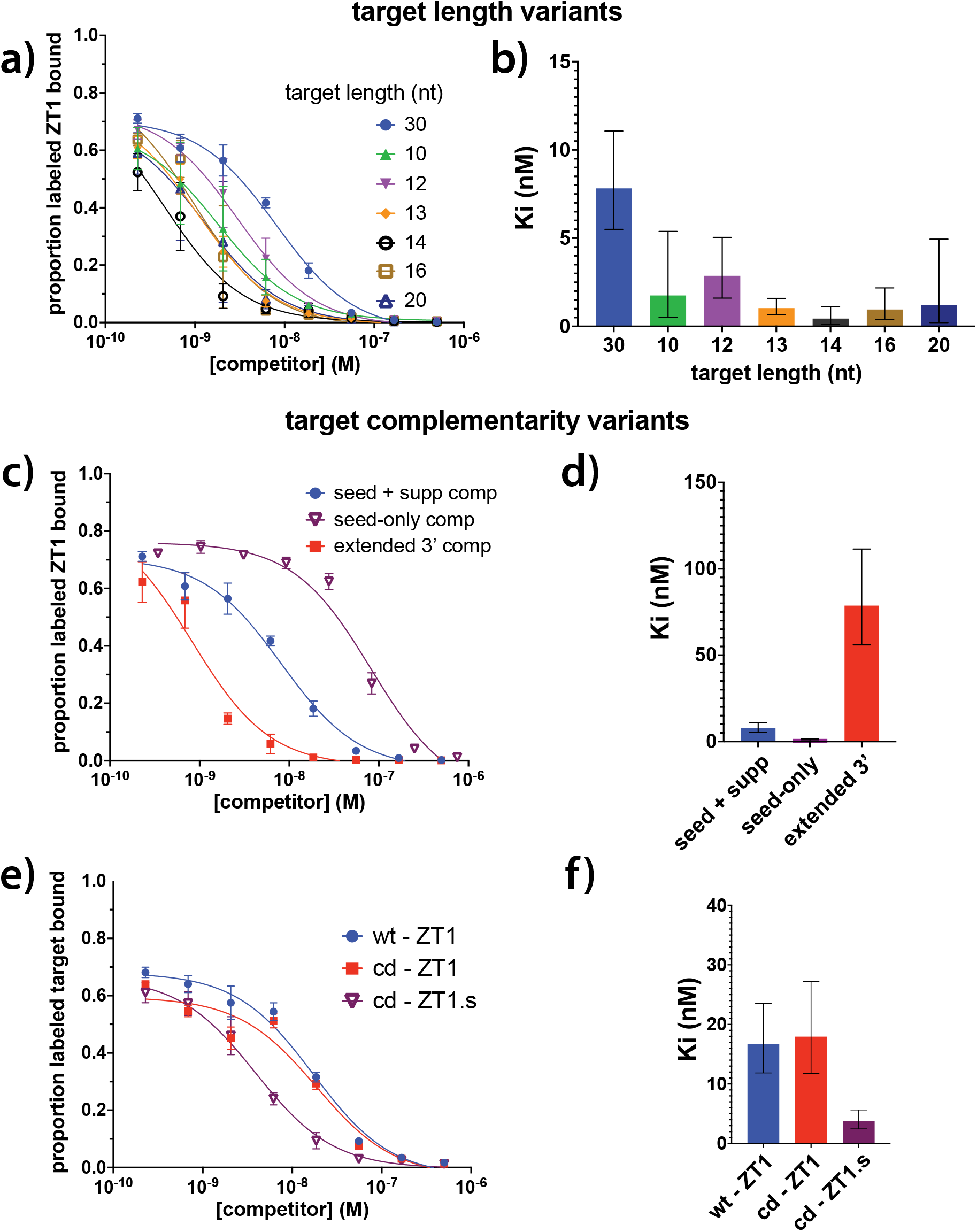
Differences between the amounts of phosphorylation triggered by different targets cannot be explained by differences in target binding. Competition filter-binding assays of target length (a-b) and complementarity (c-e) variants used were carried out in triplicate. Curves are shown on the left with 95% confidence interval (CI) and the calculated Ki for each target is shown on the right, ± SE. cd, catalytically-dead; ZT1.s, sliceable (fully-complementary) ZT1-based target.

**Figure – figure supplement 2:**
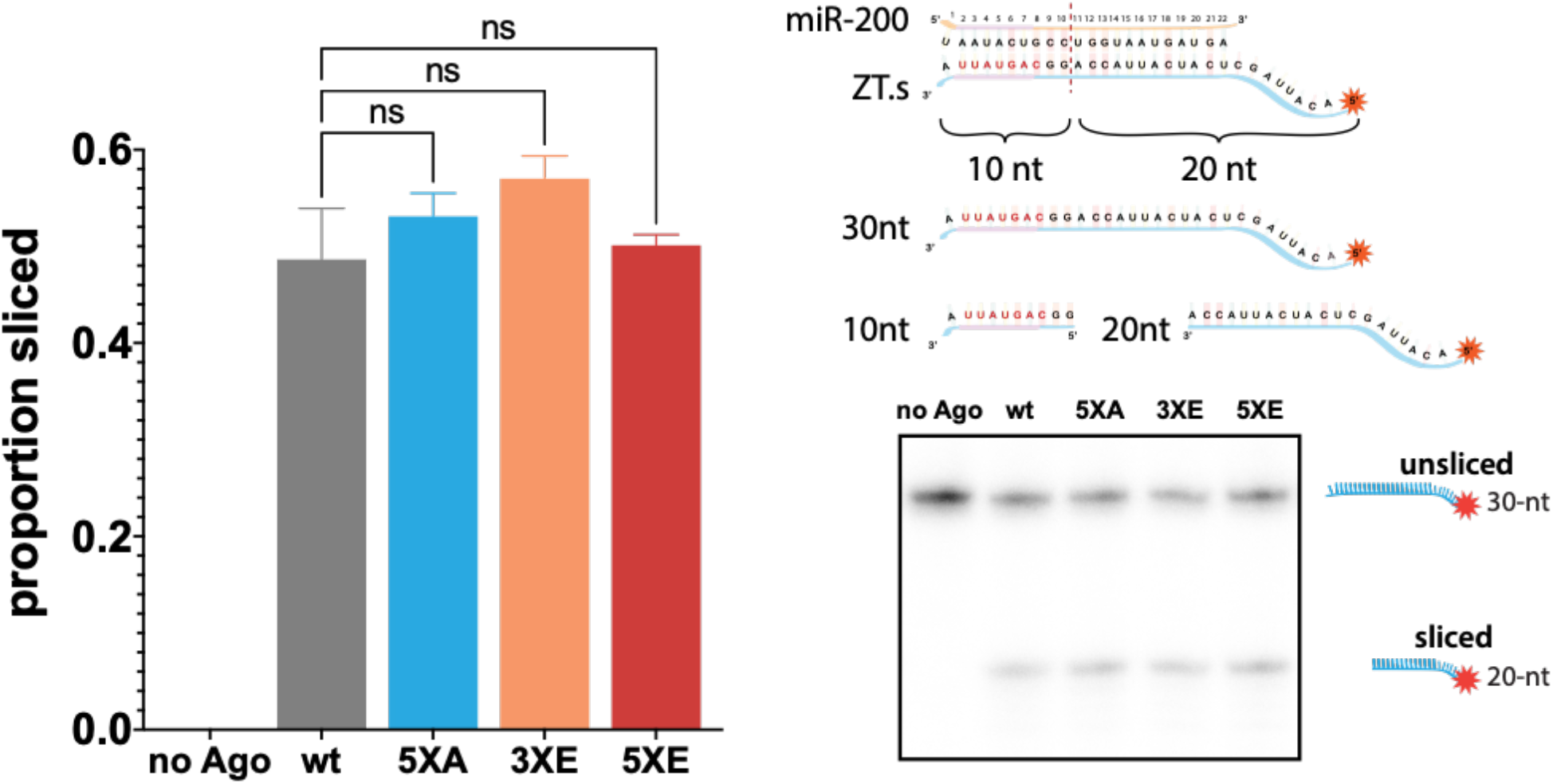
Phosphomimicking doesn’t impact slicing. The ability of various hAgo2 EI to slice a perfectly-complementary ZT1 target (ZT1.s) were tested with a slicer assay. Reactions were performed in triplicate and a representative urea PAGE gel is shown on the right with proportion sliced graphed on the left ± SE. ns, non-significant.

**Figure 3 – figure supplement 1:**
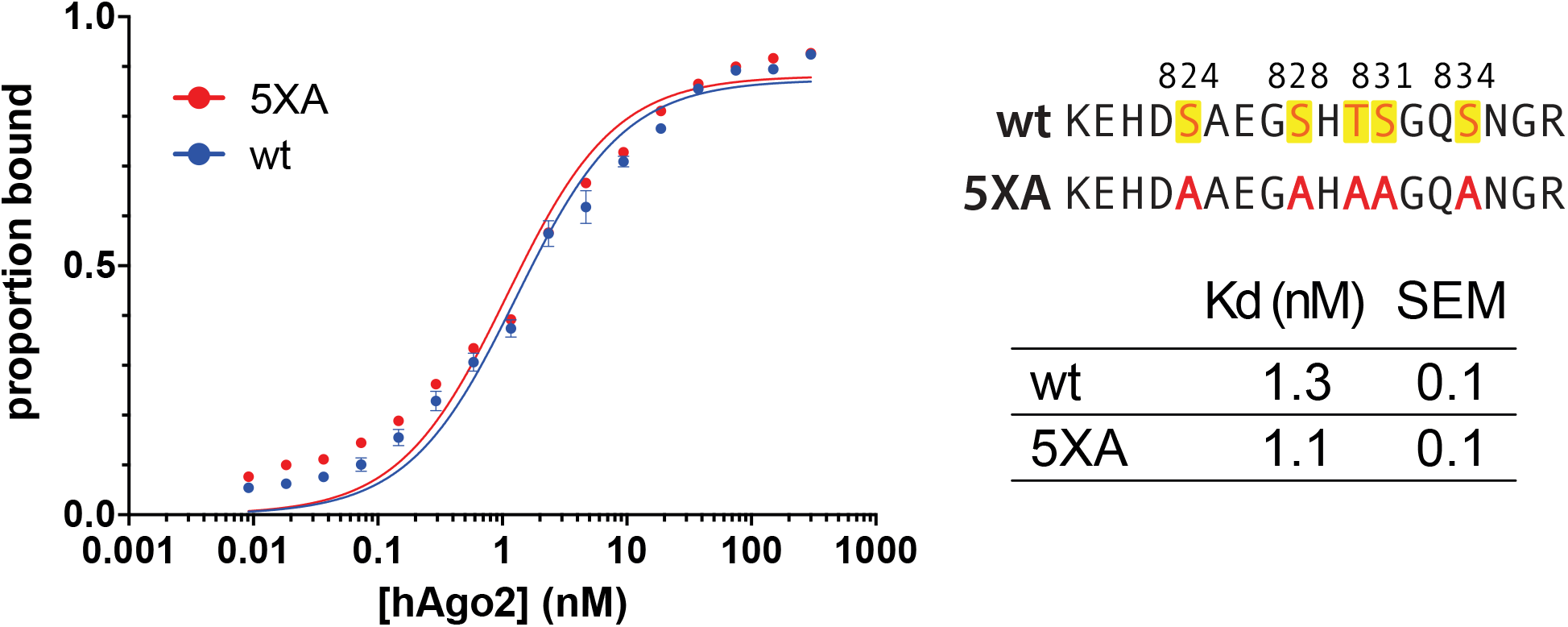
5XA substitution does not alter hAgo2’s target affinity. Filter binding assays of miR-200-bound wild-type (wt) and 5XA hAgo2 with ZT1 target were carried out in triplicate. Curves are shown on the left ± SE and the calculated Kd for each target is shown on the right. Kds were determined through non-linear regression using GraphPad Prism, one-site specific binding.

**Figure 3 – figure supplement 2:**
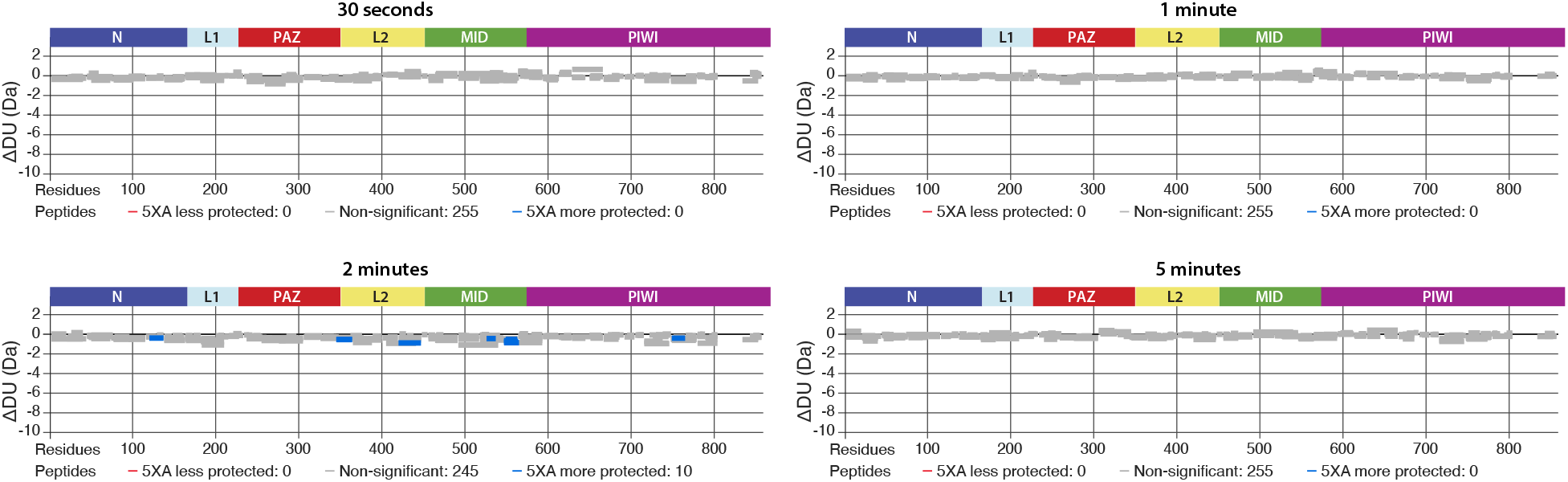
5XA and wt hAgo2 show near-identical deuterium exchange patterns, including in regions surrounding the EI, indicating that 5XA can serve as a proxy for the wt, for which we were unable to collect reliable EI coverage. Wood’s Differential plots comparing the absolute deuterium uptake of RNA-free wt hAgo2 and RNA-free 5XA hAgo2 at each time point. Statistically-different (p <0.1) peptides are highlighted with blue indicating that the 5XA is more protected than the wt. Positions of hAgo2 domains are displayed above. There were no statistically-different peptides at 30s, 1 min, or 5 min. Very minor, yet statistically-significant (p<0.01), differences were seen at the 2 min time point and were restricted to 10 (out of 255) peptides, none of which were in the region surrounding the EI.

**Figure 3 – figure supplement 3:**
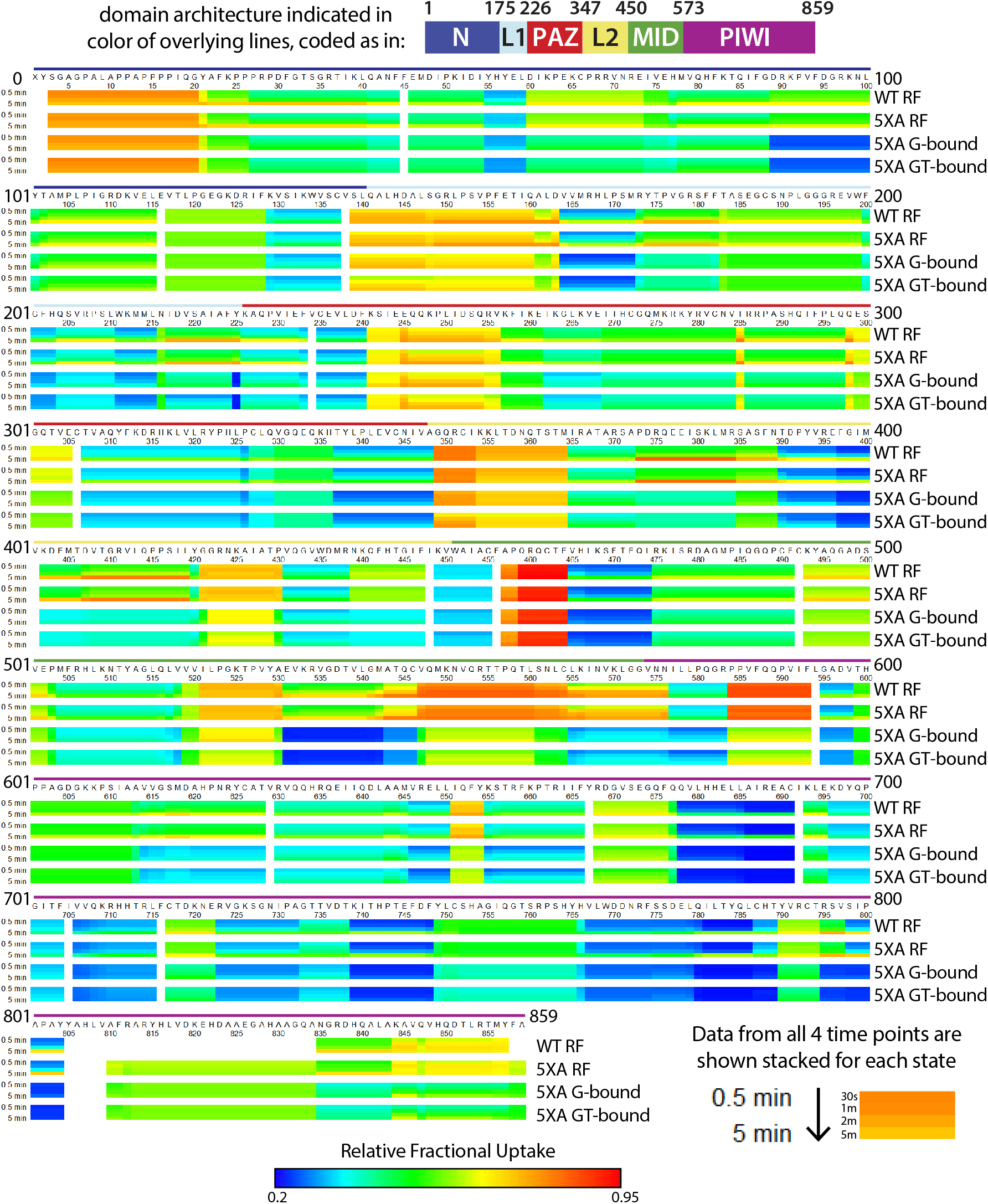
Heat map showing relative deuterium uptake for wild-type (wt) hAgo2 in RNA-free (RF) state, and 5XA hAgo2 in RF, guide-bound (G-bound), and guide+target-bound (GT-bound) states across the entire protein and all time points. Data for all four time points (30 s, 1, 2, and 5 min) are stacked and colored on a rainbow spectrum with dark blue indicating least deuterium uptake (most-protected regions) and dark red indicating highest levels of deuterium uptake (least-protected regions). The color of the overlying lines indicates the corresponding domain.

**Figure 4 – figure supplement 1:**
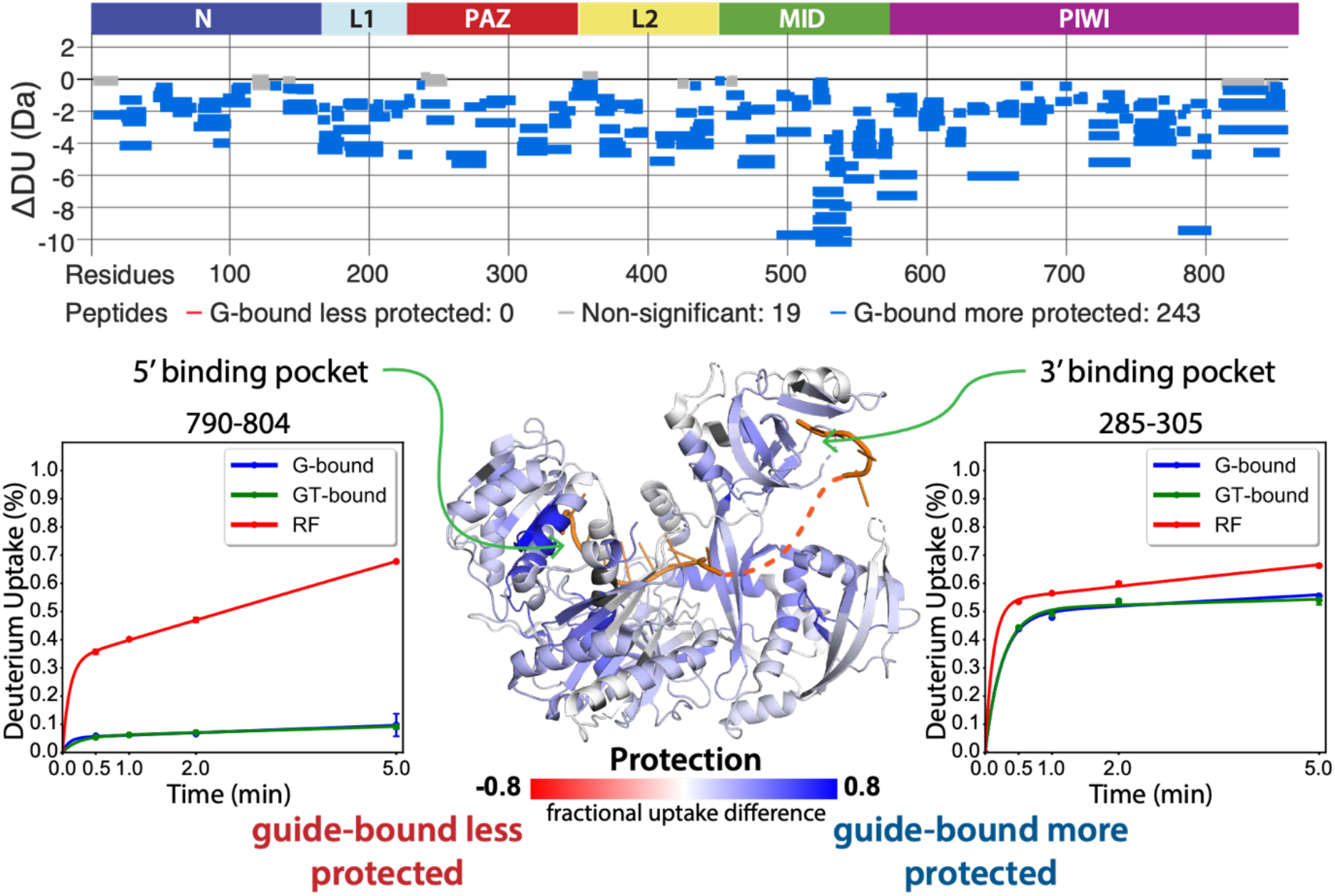
Guide-binding offers strong protection from deuterium exchange over most of hAgo2. top: Wood’s Differential plot comparing the absolute deuterium uptake of RNA-free 5XA hAgo2(RF) and guide-bound 5XA hAgo2 (G-bound) at the 5-minute time point. Statistically-different (p <0.1) peptides are highlighted in blue indicating that the guide-bound is more protected than the RNA-free. Positions of hAgo2 domains are displayed above. Bottom middle: Fractional uptake difference comparing guide-bound and RNA-free hAgo2 at the 5-minute time point. Statistically-different peptides were used to generate a heat map displayed on the structure of hAgo2 in complex with miR-20 (PDB code 4F3T, Elkayam et al. (2012)). Regions lacking HDX-MS coverage are colored dark gray in this and similar figures. Insets show deuterium uptake plots for peptides in the 5’-binding pocket (left) and 3’-binding pocket (right) showing protection offered upon guide binding.

**Figure 5 – figure supplement 1:**
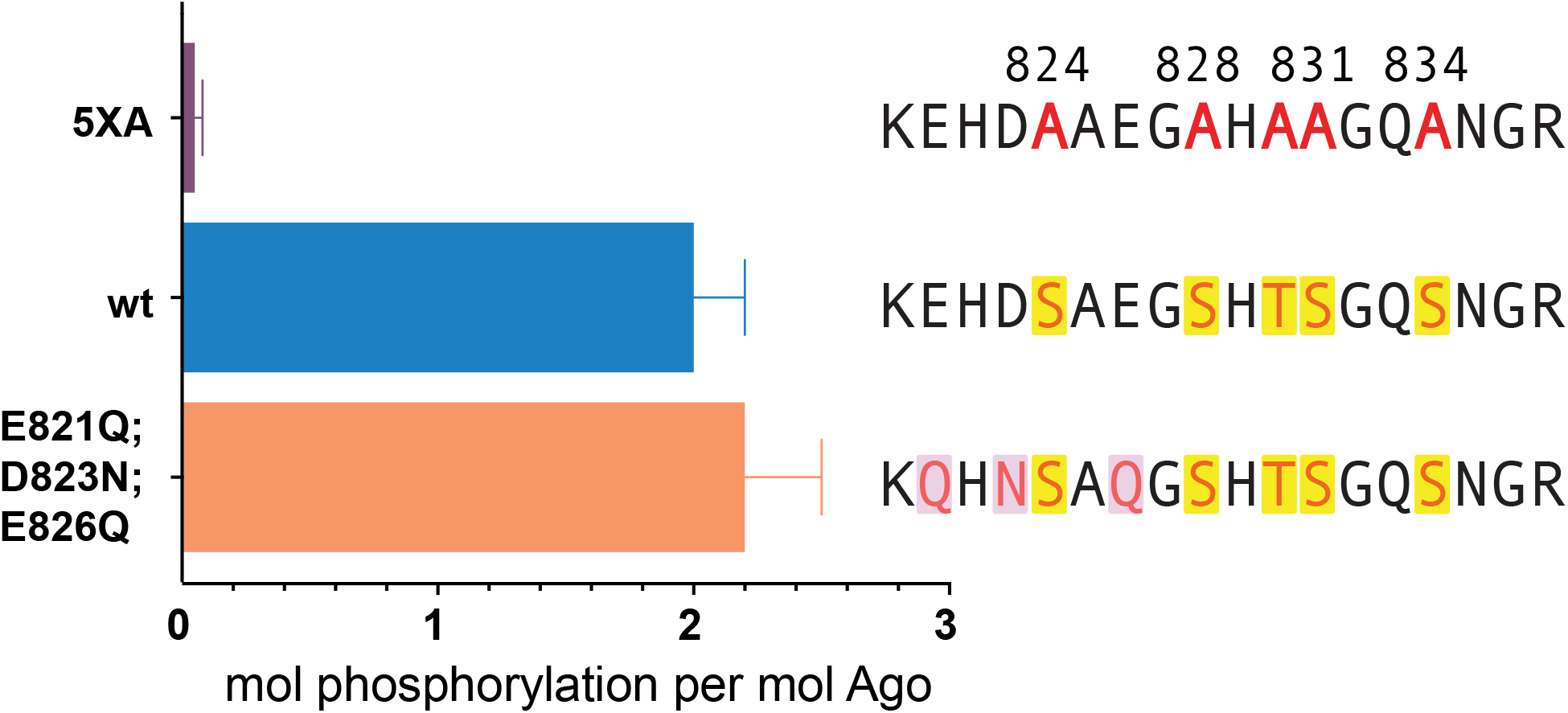
The cluster of upstream acidic residues does not play a priming role. In vitro phosphorylation assays show no effect on measured phosphorylation of hAgo2 when the residues proposed to play a priming role are neutralized, hAgo2(E821Q/D823N/E826Q). Quantification of phosphorylation was done based on spotted ATP and plotted ± SE as described in Materials and Methods. n=3

**Figure 5 – figure supplement 2:**
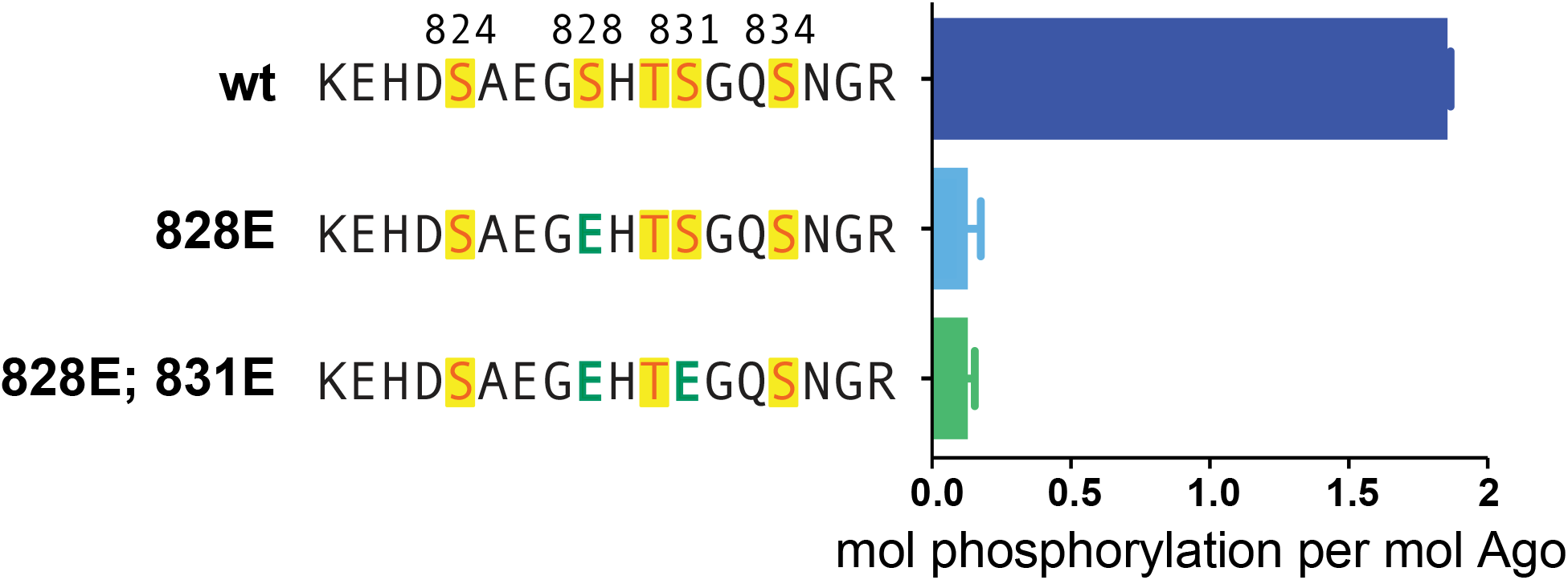
Phosphomimetics failed to prime for phosphorylation. In vitro phosphorylation assays show lack of phosphorylation of hAgo2 when serine 828 is mutated to a glutamate, with or without additional mutation of serine 831 to glutamate. Quantification of phosphorylation was done based on spotted ATP and plotted ± SE as described in Materials and Methods. n=3

**Figure 6 - figure supplement 1:**
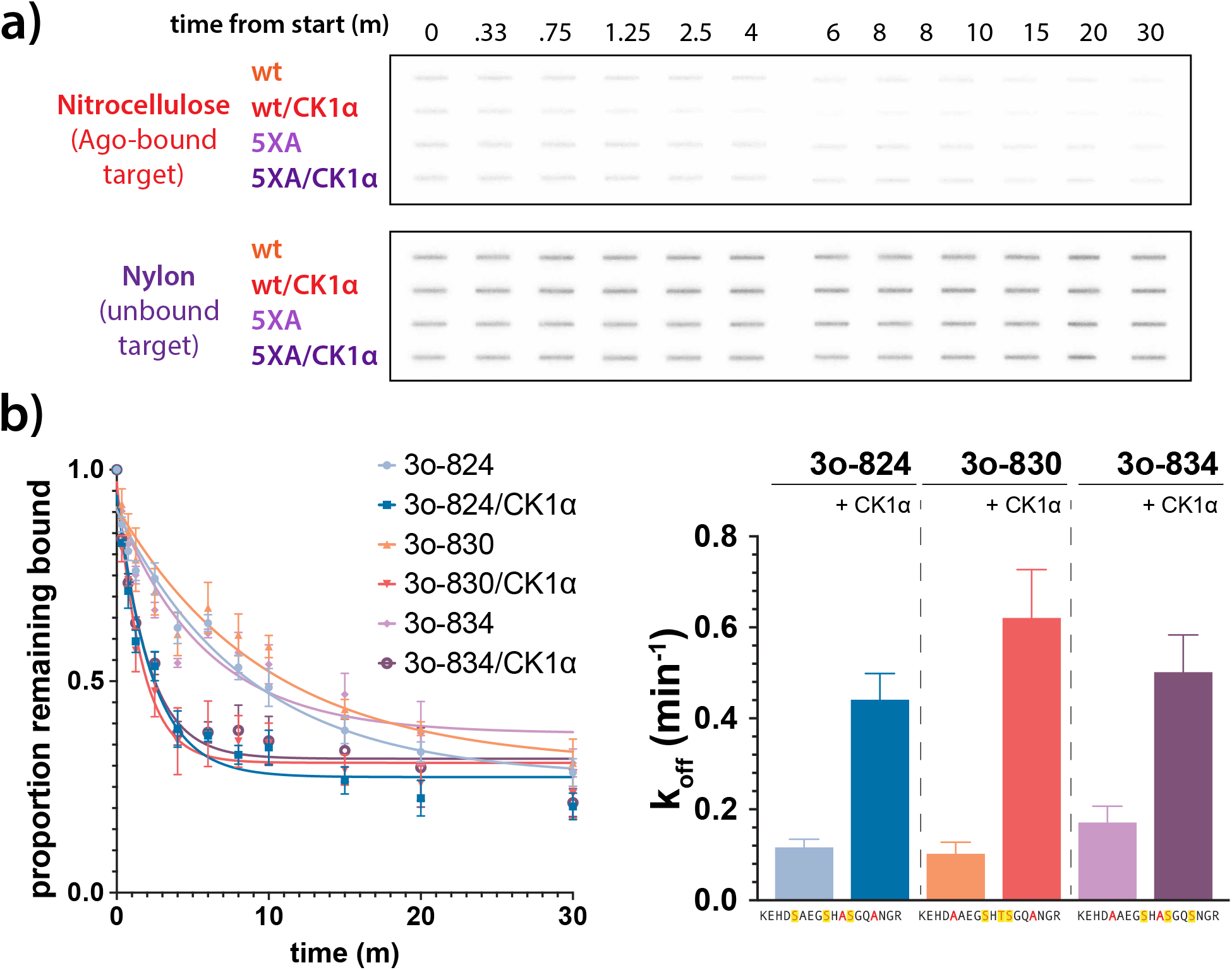
a) Representative slot bolts from target release assays showing the decrease in target bound to hAgo2 (captured on the top, nitrocellulose membrane) over time in the absence or presence of phosphorylation. b) Different triple open combinations show similar decreases in half life in response to phosphorylation, suggesting that overall quantity of phosphorylation, rather than its location, is important for promoting target release. Proportion of target remaining bound over time since addition of an unlabeled target chase is plotted with 95%CI (left) and half lives calculated using a One-Phase Exponential Decay equation in GraphPad Prism are shown ± SE (right). 3o-824, hAgo2(830A, 831A); 3o-830, hAgo2(824A, 831A); 3o-831, hAgo2(824A, 830A).

**Supplemental table 1:**
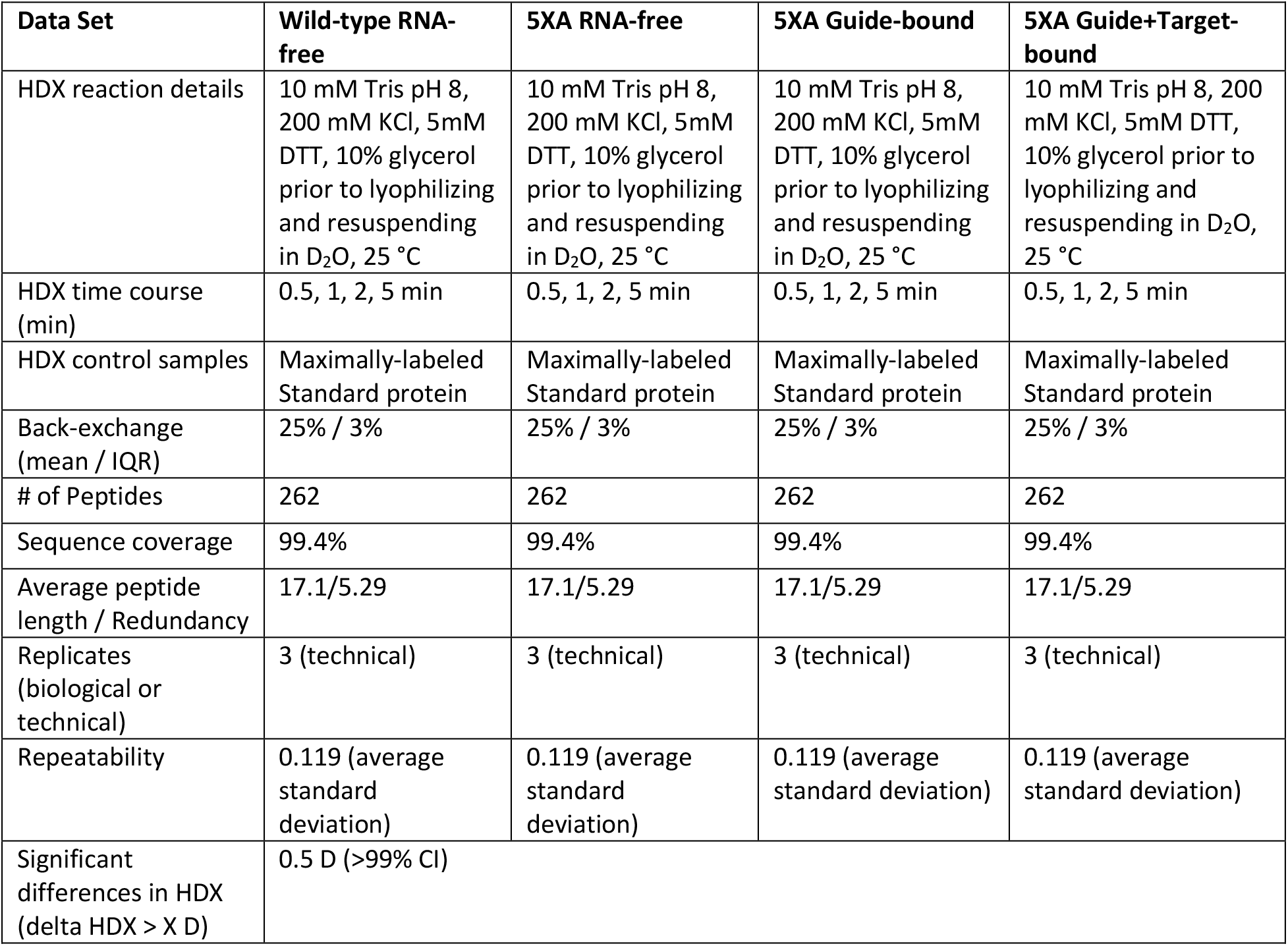
HDX Summary Data.

| **Data Set** | **Wild-type RNA-free** | **5XA RNA-free** | **5XA Guide-bound** | **5XA Guide+Target-bound** |
| --- | --- | --- | --- | --- |
| HDX reaction details | 10 mM Tris pH 8, 200 mM KCl, 5mM DTT, 10% glycerol prior to lyophilizing and resuspending in D<sub>2</sub>O, 25 °C | 10 mM Tris pH 8, 200 mM KCl, 5mM DTT, 10% glycerol prior to lyophilizing and resuspending in D<sub>2</sub>O, 25 °C | 10 mM Tris pH 8, 200 mM KCl, 5mM DTT, 10% glycerol prior to lyophilizing and resuspending in D<sub>2</sub>O, 25 °C | 10 mM Tris pH 8, 200 mM KCl, 5mM DTT, 10% glycerol prior to lyophilizing and resuspending in D<sub>2</sub>O, 25 °C |
| HDX time course (min) | 0.5, 1, 2, 5 min | 0.5, 1, 2, 5 min | 0.5, 1, 2, 5 min | 0.5, 1, 2, 5 min |
| HDX control samples | Maximally-labeled Standard protein | Maximally-labeled Standard protein | Maximally-labeled Standard protein | Maximally-labeled Standard protein |
| Back-exchange (mean / IQR) | 25% / 3% | 25% / 3% | 25% / 3% | 25% / 3% |
| # of Peptides | 262 | 262 | 262 | 262 |
| Sequence coverage | 99.4% | 99.4% | 99.4% | 99.4% |
| Average peptide length / Redundancy | 17.1/5.29 | 17.1/5.29 | 17.1/5.29 | 17.1/5.29 |
| Replicates (biological or technical) | 3 (technical) | 3 (technical) | 3 (technical) | 3 (technical) |
| Repeatability | 0.119 (average standard deviation) | 0.119 (average standard deviation) | 0.119 (average standard deviation) | 0.119 (average standard deviation) |
| Significant differences in HDX (delta HDX > X D) | 0.5 D (>99% CI) |  |  |  |

